# Multi-*e*GO: an *in-silico* lens to look into protein aggregation kinetics at atomic resolution

**DOI:** 10.1101/2022.02.18.481033

**Authors:** Emanuele Scalone, Luca Broggini, Cristina Visentin, Davide Erba, Fran Bačić Toplek, Kaliroi Peqini, Sara Pellegrino, Stefano Ricagno, Cristina Paissoni, Carlo Camilloni

**Affiliations:** Dipartimento di Bioscienze, Università degli Studi di Milano, Milano, Italy; Institute of Molecular and Translational Cardiology, IRCCS Policlinico San Donato, San Donato Milanese, Italy; DISFARM, Dipartimento di Scienze Farmaceutiche, Sezione Chimica Generale e Organica, Università degli Studi di Milano, Milano, Italy

**Keywords:** Protein aggregation, Molecular dynamics, Aggregation kinetics, Structure based models

## Abstract

Protein aggregation into amyloid fibrils is the archetype of aberrant biomolecular self-assembly processes, with more than 50 diseases associated that are mostly uncurable. Understanding aggregation mechanisms is thus of fundamental importance and goes in parallel with the characterization of the structures of the transient oligomers formed in the process. Oligomers have been proven elusive to high-resolution structural techniques, while the large sizes and long-time scales typical of aggregation processes have limited, so far, the use of computational methods. To surmount these limitations, we introduce here multi-*e*GO, an atomistic, hybrid structure-based model, which leveraging on the knowledge of monomers conformational dynamics and of fibril structures, can efficiently capture the essential structural and kinetics aspects of protein aggregation. Multi-*e*GO molecular dynamics simulations can describe the aggregation kinetics of thousands of monomers. The concentration dependence of the simulated kinetics, as well as the structural features of the resulting fibrils, are in qualitative agreement with *in vitro* experiments on an amyloidogenic peptide of Transthyretin, a protein responsible for one of the most common cardiac amyloidosis. Multi-*e*GO simulations allow to observe in time and at atomic resolution the formation of primary nuclei in a sea of transient lower order oligomers, to follow their growth and the subsequent secondary nucleation events, till the maturation of multiple fibrils. Multi-*e*GO, combined with the many experimental techniques deployed to study protein aggregation, can provide the structural basis needed to advance the design of molecules targeting amyloidogenic diseases.

**Significance Statement:** Alzheimer’s and Parkinson’s diseases are uncurable pathologies associated to the aberrant aggregation of specific proteins into amyloid fibrils. Understanding the mechanism leading to protein aggregation, by characterizing the structures of the oligomeric species populated in the process, would have a tremendous impact on the design of therapeutic molecules. We propose that a structure-based approach to molecular dynamics simulations can allow following at high resolution the aggregation kinetics of thousands of monomers. Having shown that simulations can describe the aggregation of a Transthyretin amyloidogenic peptide, we demonstrate how their efficiency allows acquiring a wealth of structural information. We foresee that integrating the latter with the many techniques developed to study protein aggregation will support the design of molecules to modulate amyloidogenesis.

## Introduction

Amyloid fibril formation is a highly specific self-assembly process, requiring a large degree of similarity between the interacting amino acid sequences (1). Amyloids, resulting from the uncontrolled transition of normally soluble proteins, were originally found in association with neurodegenerative diseases (2, 3) only more recently they have been also associated to several physiologic functions (4, 5). Amyloid fibrils share a cross-β architecture. β-strands are oriented perpendicularly to the fibril axis allowing the formation of a dense network of intermolecular hydrogen bonds. Sidechains, instead, contribute to both intra and inter-molecular interactions (6, 7). *In vitro*, the amyloid fold seems to be accessible to a large number, if not all, proteins (ordered or disordered) or even short sequences of amino acids (8, 9). Thermodynamic considerations, indeed, suggest that native proteins are metastable species in physiological conditions, with the global free-energy minimum corresponding to their amyloidogenic state (10).

Protein aggregation into amyloid fibrils is inherently a dynamic process. Many interconverting species of differing sizes and structures can be populated over multiple time scales (11). The description of amyloid fibrils formation thus requires understanding not only the properties of the end states, *i*.*e*., monomers and fibrils, but also of the different oligomeric species transiently populated in between. Remarkably, in diseases like Parkinson’s, Alzheimer’s, type 2 diabetes mellitus and cardiac amyloidosis, some oligomeric species may be the primary pathogenic agents (12–16). Furthermore, toxic oligomers have been found in model proteins and associated to specific physico-chemical properties like size and hydrophobicity, even if it is not yet clear if these are relevant for all amyloidogenic diseases (17–19). Structural approaches based on solid-state nuclear magnetic resonance (ssNMR) and cryogenic electron microscopy (cryo-EM) are revealing the atomic structure of amyloid fibrils formed by different proteins in diverse conditions (6, 7, 20– 22). The aggregation process itself can be only studied at very low resolution, by aggregation kinetics assay, where experimental conditions are tuned to let a solution of monomeric protein interconvert into amyloid fibrils. Seeds obtained by previously formed fibrils can also be employed to catalyze the interconversion (23).

Chemical kinetics analysis provides a framework to dissect the microscopic mechanisms at play in fibril formation (24). Aggregation is described by a network of microscopic processes like primary nucleation and elongation, as well as secondary nucleation processes, like fragmentation and surface induced nucleation. By globally fitting multiple accurate aggregation kinetic traces obtained for multiple initial monomer concentration it is possible to estimate the rates for the different microscopic processes and use these to interpret the macroscopic observations. Such analysis highlighted how proteins associated with amyloidogenic diseases seem to prevalently aggregate through secondary nucleation mechanisms while physiological amyloids may be mainly controlled by primary nucleation (25, 26). Drug design strategies are being implemented based on the kinetic modulation of such mechanisms (27). Nonetheless, despite its power, chemical kinetics fails to provide detailed structural information of the species at play along the process.

Given the inherently transient and dynamic nature of the species populated in an assembly process, molecular dynamics (MD) simulations naturally complement current experimental approaches (27). Simulations of the aggregation kinetics have been mainly employed to characterize the early events in the oligomerization of few peptides at high concentrations because of the combination of challenges resulting from system sizes and relevant time scales. Implicit solvent models have been employed to mitigate these problems and allowed studying the oligomerization of 20 monomers of Aβ40 and Aβ42 for hundreds of *ns* (28). Simulating larger systems on longer time scales requires coarse grain models (CG). Notably, fibrils may easily be formed by tens of thousands of monomers.

CG simulations have been important to shed light on the role played by kinetics, thermodynamics, and other physico-chemical properties in protein aggregation, but they have been employed only for very qualitative studies (29–31). In the field of protein folding simulations, the most used CG models are structure-based (SB), also known as Gō models (32–34). SB models are an implementation of the principle of minimal frustration (or the folding funnel (35)): attractive interactions are defined only between amino acids or atoms close in space in the native crystal state, consequently the minimum energy configuration is the native crystal configuration. This allows studying very efficiently folding and unfolding transitions by dramatically decreasing the cost to evaluate interactions and speeding up the overall diffusion in conformational space, *e*.*g*., the folding time of a protein can be rescaled from milliseconds to hundreds of nanoseconds.

In keeping with the observation that the amyloid structure is the global free-energy minimum of a protein at high concentration (10), here we introduce multi-*e*GO, a novel hybrid SB model including non-bonded interactions derived from both the dynamics of the soluble protein as well as from the structure of the amyloid fibril, and transferable bonded interactions optimized to reproduce the results of state-of-the-art explicit solvent molecular force fields. While SB models including more than one reference structure have been employed to study large conformational changes in proteins as well as metamorphic proteins (33, 36–40), here we show that multi-*e*GO can be used to follow at high resolution the aggregation of thousands of monomers as a function of their initial concentration. Our results are qualitatively in agreement with experiments and enable the structural investigation of the aggregation of proteins into amyloid fibrils.

## Results

To develop multi-*e*GO, we used the Transthyretin 105-115 amyloidogenic peptide (TTR_105-115_) (41, 42). Transthyretin is a well-studied amyloidogenic protein responsible for both sporadic and genetic cardiac and systemic amyloidosis (43). TTR_105-115_ has been often used as a model system to study aggregation and three amyloid polymorphisms were determined at atomic resolution by a combination of multiple techniques including ssNMR and cryo-EM (20). NMR analysis of monomeric TTR_105-115_ in solution indicates that it populates primarily random-coil structure with a low percentage of turns or helical elements (44). Multi-*e*GO is built including information from the structure or the dynamics of the end states and uses them to infer the properties of the intermediate oligomeric states (**Figure S1**). To have a realistic reference conformational ensemble of monomeric TTR_105-115_ we performed an explicit solvent MD simulation using the AMBER99SBDisp force-field (45). This simulation well represented the behavior of TTR_105-115_ in solution, showing a broad flexibility and sporadic turns, as reported by the radius of gyration distribution and the per-residue contact probability map in **Figure 1**, and overall agrees with previous NMR studies (44).

**Figure 1.**
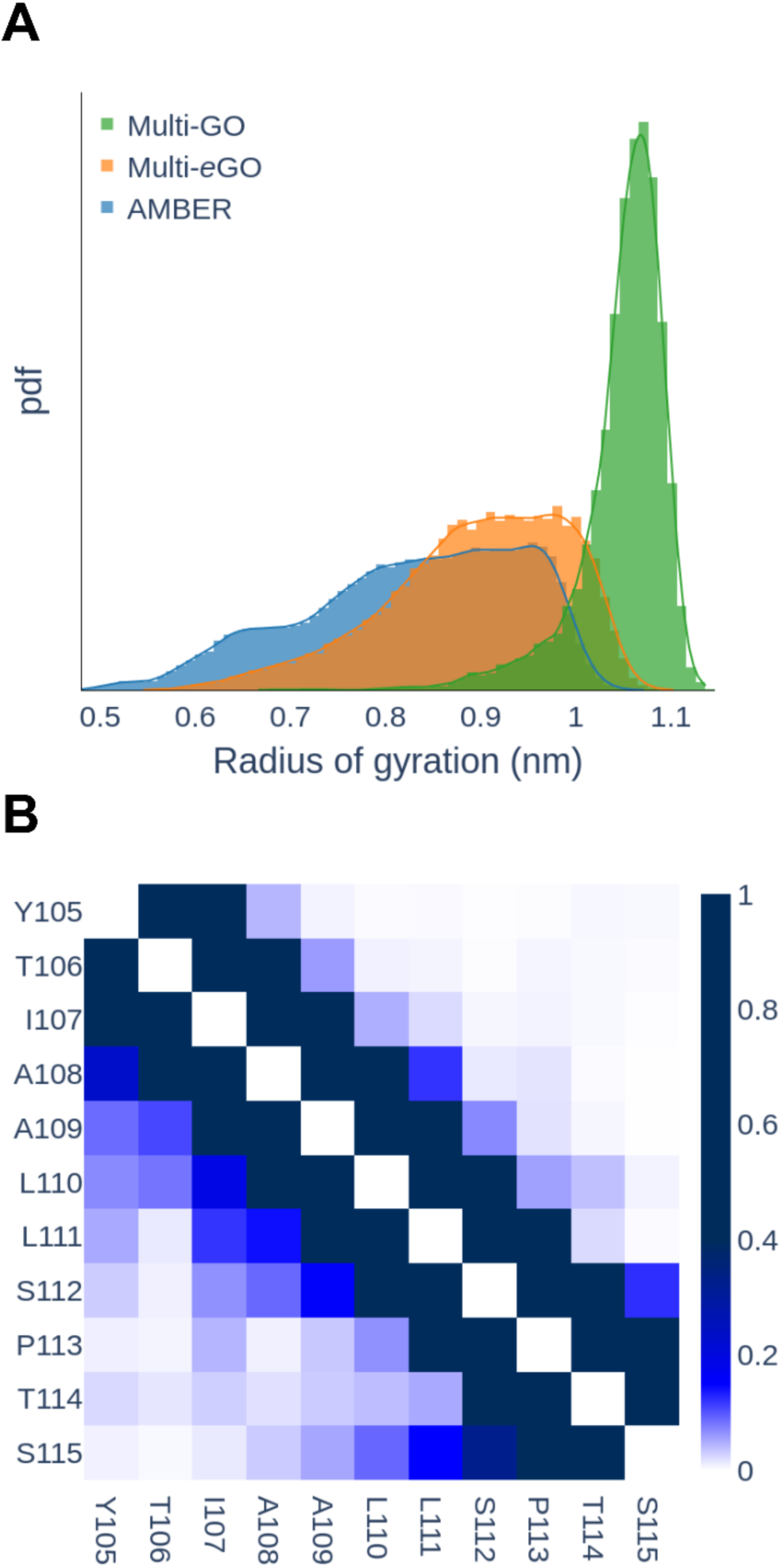
TTR_105-115_ peptide monomer dynamics. (A) Gyration radius distribution of TTR_105-115_ peptide conformational ensemble according to multi-GO (green), multi-*e*GO (orange), and AMBER (blue) simulations, respectively. The multi-GO distribution describes an open conformation with a single peak at 1.07 nm. The AMBER simulation shows multiple peaks over a broad range of values. The multi-*e*GO distribution is shifted towards more extended conformations than AMBER but still shows a broad range of values in qualitative agreement with the former. (B) Per residue probability contact map for the AMBER (lower left) and multi-*e*GO (upper right) simulations.

### Multi-eGO reproduces the conformational dynamics of TTR_105-115_ in solution

Following previous works on metamorphic proteins (38, 39), we first defined the multi-GO SB force field, at all heavy atom (non-hydrogens) resolution, as a combination of terms obtained from two reference structures (cf. Material and Methods), namely the protein in its native monomeric state (extracting TTR_105-115_ coordinates from pdb 4TLT) and the amyloid fibril (pdb 2M5K), cf. **Figure S1** in the Supplementary Information. A Multi-GO simulation of a TTR_105-115_ monomer explored only extended configurations with an average radius of gyration of 1.05 nm, in comparison with 0.83 nm of the AMBER one (**Figure 1A**) and the conformational ensemble did not show long range contacts, cf. **Figure S2** in the Supplementary Information. The multi-GO ensemble described above did not capture the conformational freedom of the monomeric state, and consequently may not capture that of early intermediate oligomeric states.

To increase the descriptive power of the model we introduced multi-*e*GO as a hybrid transferable/SB model (cf. Materials and Methods). The most relevant differences are that all bonded interactions, and in particular proper dihedral angles, are transferable, while non-bonded interactions are learned from a reference simulation for the monomeric state, i.e., the AMBER force-field simulation introduced above, and a reference amyloid fibril structure (pdb 2M5K). Remarkably, while the multi-GO simulation explored only extended configurations, the multi-*e*GO model could better recapitulate TTR_105-115_ dynamics in solution with an average radius of gyration of 0.90 nm. The contact probability map for AMBER and multi-*e*GO, showed in **Figure 1B**, indicate that multi-*e*GO can also qualitatively describe the intermolecular transient interactions of the peptide.

### Multi-eGO can simulate TTR_105-115_ aggregation

Using multi-*e*GO, a total of 15 simulations (each involving 4000 TTR_105-115_ monomers) as a triplicate of five different concentrations, between 7 and 13 mM, were produced. The resulting aggregation kinetics are shown in **Figure 2A** as the number of monomers forming assemblies, from decamers to larger ones, at a given time. Of note, the time scale of the simulations is only nominal and comparison with experiments should consider a scaling factor. The simulations displayed sigmoidal concentration dependent kinetics, where an increase in monomer concentration resulted in a reduction of the lag phase. We also observed that the variability of the curves increased inversely with the initial monomer concentration (46). From the resulting curves we obtained the half time, *τ*_1/2_, showed in **Figure 2B**, and the growth rate, *r*, as the slope of the straight line fitting the region of the curve around *τ*_1/2_ (**Figure S3** in the Supplementary Information). The double log plot of *τ*_1/2_ as a function of the concentration, **Figure 2B**, showed a bilinear trend with a change of slope, the scaling exponent *γ*, at concentrations lower that 8.5 mM, suggesting that at high monomer concentration a dominant aggregation mechanism becomes saturated (47).

**Figure 2.**
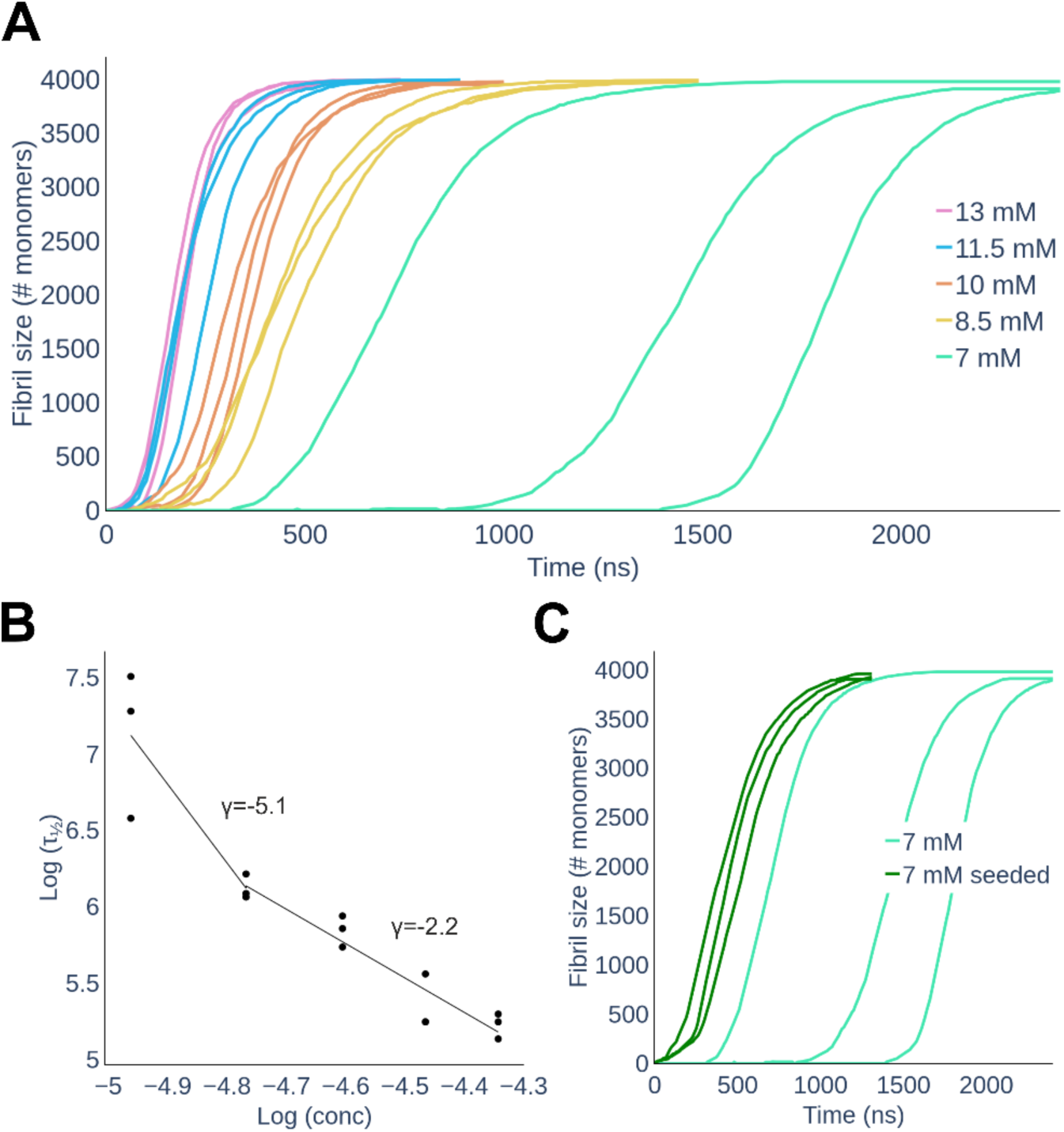
(A) Simulated aggregation kinetics. Curves represent the number of monomers involved in an aggregate of at least 10 monomers as a function of nominal simulation time. (B) Log-log plot of the half times, *τ*_1/2_, as a function of the initial monomer concentration. The points are fitted with two straight lines in the range 7 to 8.5 mM and 8.5 to 13 mM. (C) Aggregation kinetics of seeded and unseeded 7 mM simulations. Curves represent the number of monomers involved in an aggregate of at least 10 monomers as a function of nominal simulation time. The addition of the seeds reduces *τ*_1/2_ and makes it less variable compared to the unseeded simulations, leaving the slope of the growth is unaffected.

To test the ability of multi-*e*GO to capture differences between seeded and unseeded aggregation kinetics, we also performed three seeded simulations at 7 mM by adding as seed a 10 monomers oligomer obtained from previous 13 mM simulations (cf. *Structural details of TTR*_*105-115*_ *aggregation kinetics*, in the following). In **Figure 2C** it is shown that the addition of the seed resulted in a marked decrease of *τ*_1/2_ as well as in a reduction of its variability. The observed growth rate *r* instead, was the same for the unseeded simulations performed at the same concentration (i.e., 7 mM).

### Multi-eGO TTR_105-115_ simulations can form polymorphic fibrils

The eighteen simulations performed at 5 different concentrations yielded a total of 41 distinct fibrils. These fibrils grew in length from 163 Å to 515 Å with an average length of 360 Å (**Table 1** and **Table S1** in the Supporting Information). All fibrils displayed the expected cross-β topology with a parallel and in-register stacking of chains in the same β-sheet as shown in **Figure 3**. The average distance between β-strands in the cross β-sheet was 4.7 Å. Facing β-sheets were antiparallel and shifted by 2.5 Å, resulting in the even-numbered side chains of one peptide interacting with the odd-numbered side chains of two opposite peptides. Following previous nomenclature (20), we define protofilament a structure made of two antiparallel β-sheets, the further addition of two β-sheets in a protofilament determines a filament (**Figure 3A**). Here, the β-sheet content in a filament could vary from 4 to 17 with an average of 10. A filament could grow in the peptide chain direction through interactions between the N- and C-terminal residues (**Figure 3B**), determining then a fibril. This head-to-tail interaction resulted from Y105 side chain interacting with both S115 carboxyl group and Y105 side chain of the facing β-sheet. The number of filaments in a fibril could vary from 2 to 6 with an average of 4. Mature fibrils displayed a twist per monomer between −0.1° and −0.85° measured as the torsion angle between two vectors obtained from Y105-Cα and S115-Cα carbons of subsequent molecules in the same β-sheet. Single filaments displayed a more pronounced twist of −5° compared to mature fibrils (**Figure 3**). At higher concentration we saw the formation of more fibrils indicating that more nuclei are produced compared to lower concentration (**Figure S4**). Since the number monomer was fixed at 4000, the fibrils grown at higher concentration were shorter in length than the one obtained at lower concentration (cf. **Table 1** and **Table S1** in the Supporting Information). We also observed, at higher concentration, fibrils adhering together (**Figure 3C**). Again, given the fixed and relatively small number of monomers, some protofilaments were not able to become fibrils due to monomers depletion. We did not observe any specific difference in the fibrils formed at 7 mM seeded and unseeded simulations.

**Table 1.**
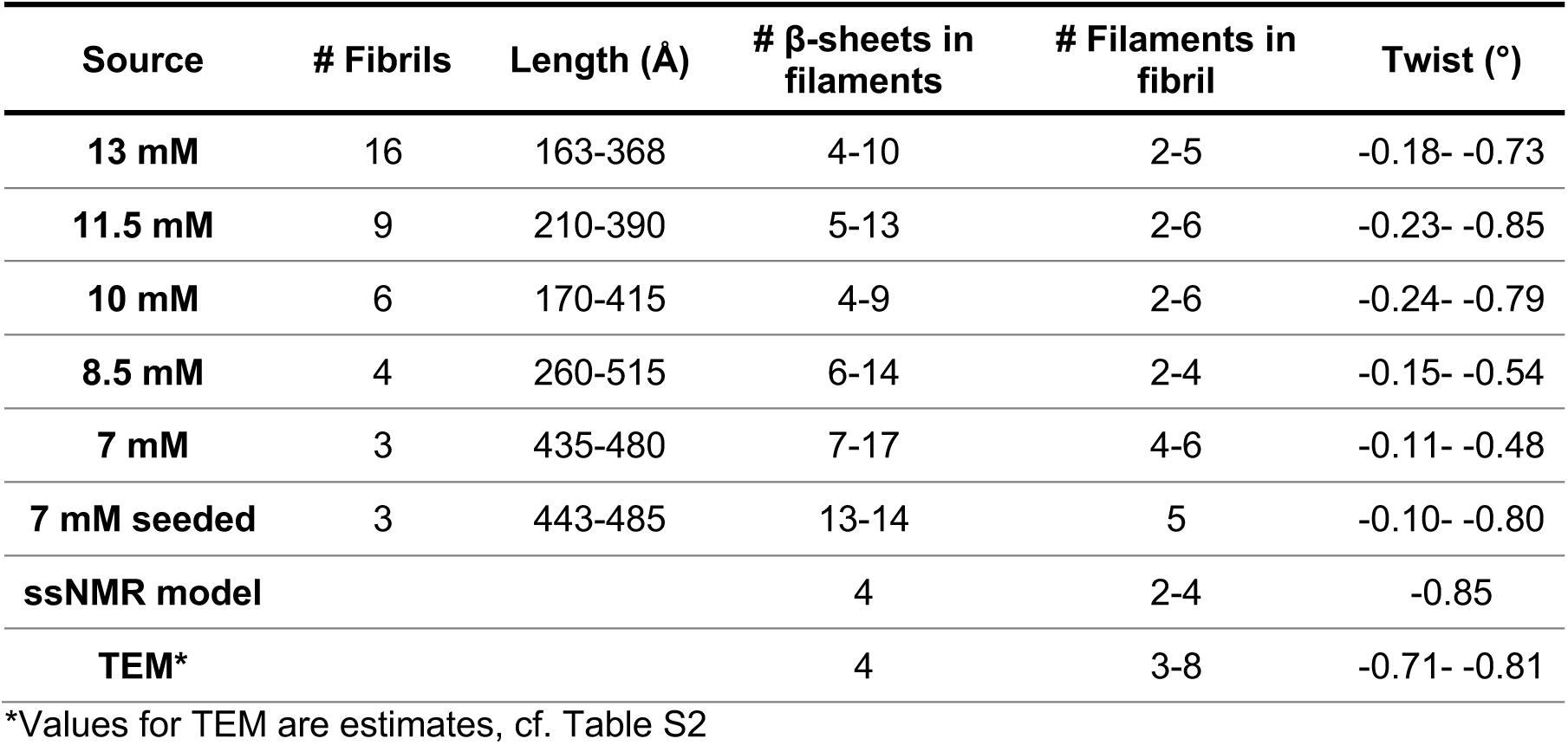
Summary of the main structural features of the fibrils discussed in this work. The first six rows indicate fibrils formed in silico in our multi-*e*GO simulations, ssNMR is for the fibrils corresponding to pdb codes 2M5K, 2M5M, and 3ZPK, and TEM are those observed in vitro in this work.

**Figure 3.**
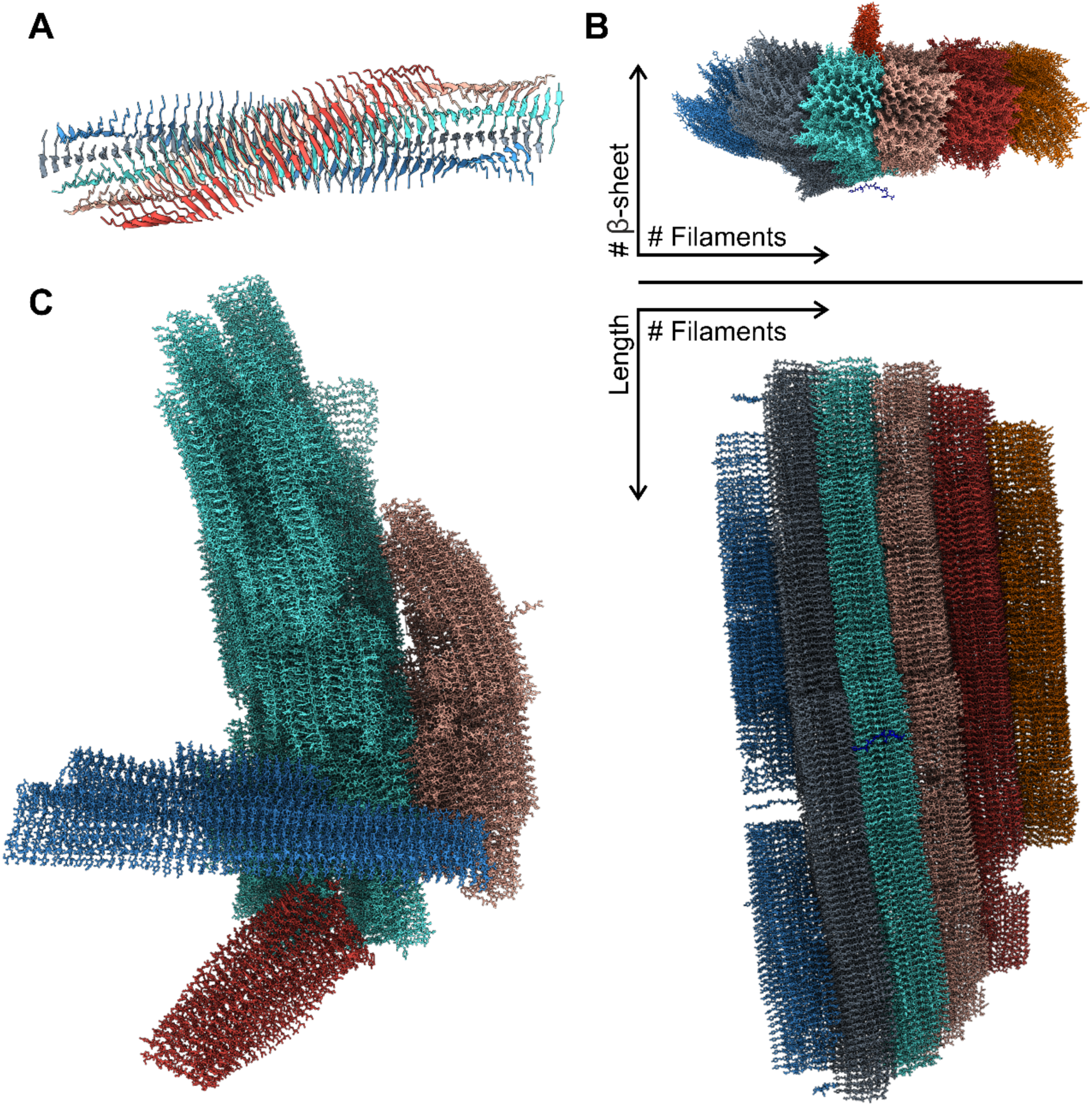
(A) A filament model observed at the end of a simulation. Colors indicate the 5 different β-sheets composing the filament. (B) Top and side view of a mature fibril with colors indicating the different filaments. From the top view It is possible to see a peptide which is about to attach to the fibril and a protofilament which is perpendicular to the main fibril. (C) Multiple mature fibrils (each represented with a different color) interacting with each other from one of the 13 mM simulations.

Compared to the reference model determined by ssNMR and Cryo-EM (20), the only remarkable difference is that our fibrils do not display any wet cavity within filaments. The cavity should accommodate structured water interacting with exposed sidechains. In our model all sidechains in a filament are tightly packed, therefore we also observed a variable number of β-sheets in a filament where the reference model contained always four of them. However, there are evidence of such variations (48, 49). The reference model described a structural polymorphism, based on the number of filaments, from doublet to quadruplet. In our simulations we saw the same polymorphism but extended to six filaments in a single fibril.

### TTR_105-115_ in vitro aggregation experiments recapitulate multi-eGO simulations

To validate the *in-silico* aggregation kinetics, we performed aggregation assays monitored by Thioflavin T (ThT) fluorescence (**Figure 4**). TTR_105-115_ peptide was incubated at 37 °C at different concentrations (i.e., 13 mM, 10 mM, and 7 mM) and ThT fluorescence was monitored for 150 h. ThT fluorescence increased over time indicating (**Figure 4A**) concentration dependent kinetics of aggregation. The lag phase at 13 mM was considerably shorter compared to the ones at 10 or 7 mM. The fluorescence plateau was reached faster in the most concentrated samples, whereas at the lowest concentration tested (i.e., 7mM) the plateau was not observed during the overall incubation time. The mean values of three independent experiments were subjected to nonlinear regression analysis, using Boltzmann sigmoidal equation. From the regression we derived experimental *τ*_1/2_ of 33.7±4.3 h, 62.0±16.6 h and 125.7±10.2 h for 13 mM, 10 mM, and 7 mM, respectively. **Figure 4B** shows in double log plot a linear correlation between peptide concentrations and half times, with a slope *γ* comparable to the one obtained from simulations in the range 8.5 to 13 mM (namely, −2.0 and −2.2 for the experiments and simulations, respectively, cf. **Figure 2B**). This is of note given the relative simplicity of our model and the fact that it does not include any specific information about the kinetics of the process.

**Figure 4.**
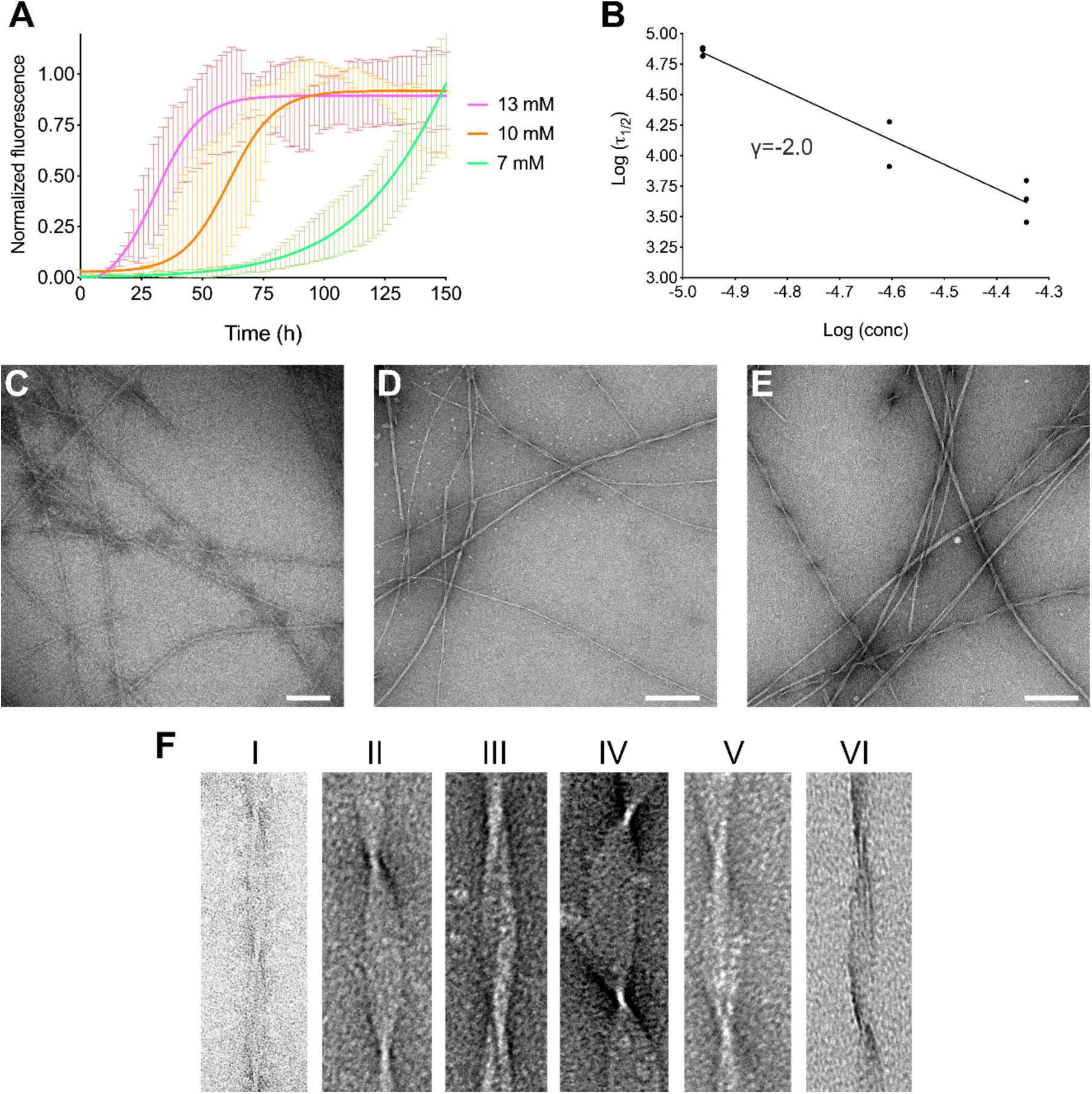
TTR peptide aggregation kinetics in vitro. (A) Aggregation kinetics of the TTR_105-115_ peptide at 13 mM, 10 mM and 7 mM are shown in magenta, orange and green, respectively. TTR peptide at 37 °C were obtained by monitoring of ThT fluorescence. The mean value of three independent experiments analyzed by linear regression using Boltzmann sigmoidal equation is reported. (B) Log-log plot of the in vitro half times, *τ*_1/2_, as a function of the initial monomer concentration. (C-E) Electron micrographs of fibrils formed by TTR_105-115_ peptide incubated at 13 mM (C), 10 mM (D) or 7 mM (E) at 37 °C for 150 h. Scale bars correspond to 100 (C) or 200 (D and E) nm. (F) Representative TEM images of the six main fibrillar morphologies. The detailed structural parameters of each morphology are reported in Table S2.

The aggregates of TTR_105-115_ peptide obtained by the aggregation kinetics experiments were negatively stained and analyzed by transmission electron microscopy (TEM). As reported in **Figure 4**, we observed remarkable polymorphism in all the conditions tested. Morphological analysis identified six main different types of structures (**Figure 4C-E** and **Table S2** in the Supporting Information). The mean width at the crossover is 37±4 Å as previously observed by Fitzpatrick et al. (20). The fibrils crossover in the six polymorphs ranges from 1041±24 Å to 1185±43 Å; the diameter varies considerably ranging from 115±11 Å to 326±21 Å. Representative pictures of each identified morphology are reported in **Figure 4F**, and the main fibril parameters in **Table S2**. Notably, the observed widths between crossovers correspond to multiples of the peptide chain length, suggesting the presence of up to 8 aligned filaments (**Table S2**).

### Multi-eGO can provide structural details for TTR_105-115_ aggregation kinetics

Having shown that multi-*e*GO could simulate the aggregation of TTR_105-115_ from monomers to fibrils with structural and kinetic features compatible with experiments, we could then look at the structures populated along the self-assembly process in detail. In **Figure 5A** and **Figure S5** in the Supporting Information, is shown the number of monomers, dimers, and trimers for our aggregation kinetics as a function of time. We observed that the number of free monomers displayed a sigmoidal behavior symmetric with respect to that of the fibril size. The number of dimers and trimers showed instead a noisy but relatively constant trend till the end of the lag phase (*t*_*lag*_), defined as the intersection between a straight-line, tangent to aggregation kinetic curve at *τ*_1/2_ with slope *r*, and the time axis, and quickly dropped after this time, suggesting that once fibrils start to grow, most monomers contributed to the fibril growth instead of forming new oligomers and the ones already formed before *t*_*lag*_ dissolve over time. In **Figure 5B** is shown the time resolved distribution of oligomer sizes of the first 13 mM simulation before *t*_*lag*_ (cf. **Figure S6** for all simulations). This analysis allows to follow the emergence and growth of primary nuclei and suggested that fibrils stem from primary nuclei made of around 10 monomers. Assuming the simulations prior to *t*_*lag*_ at equilibrium, and thus averaging over this time window, we observed how, at all concentrations, dimers and trimers were the most represented oligomeric species with populations in the 5-10% and 0.5-2% range, respectively. Higher order oligomers were scarcely populated stressing the need to simulate large numbers of monomers to study aggregation (cf. **Figure 5C**).

**Figure 5.**
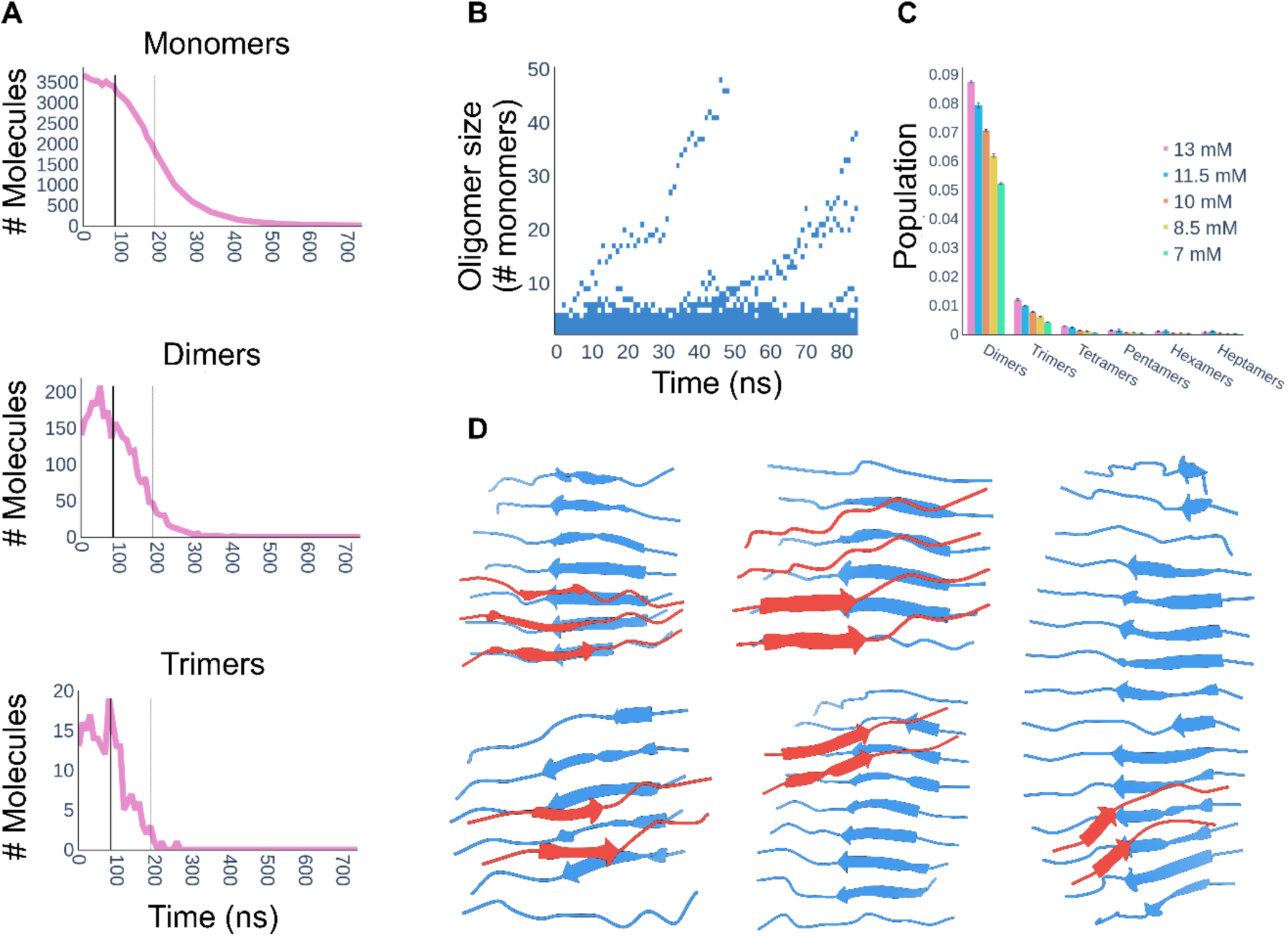
(A) Number of monomers, dimers, and trimers over time of the first 13 mM simulation. In each plot is reported t_lag_ (solid thick line) and *τ*_1/2_ (dashed thin line). The monomer decrease is symmetrical compared to fibril growth (cf. Figure 2A). The number of dimers and trimers is relatively constant till t_lag_ and quickly drops after this time. (B) Time resolved evolution of oligomers order of the first 13 mM simulation before *t*_*lag*_. (C) Oligomer order populations in the time window from 0 to *t*_*lag*_ and averaged over the three replicates. The different colors represent the different concentrations, while the error bars represent the standard deviation over the three simulations. In this time window only monomer and dimers are significantly populated (C), nonetheless one can observe the emergence of two fibrils (B). (D) Five representative structures of primary nuclei extracted from five different simulations. The β-sheets colored in blue, representing the initial oligomer, are stabilized as primary nuclei by interacting with the second β-sheet colored in red.

The structures of oligomers involved in primary nucleation are shown in **Figure 5D**. All primary nuclei displayed two antiparallel β-sheets. Observing the trajectory, we were able to describe their formation. Free monomers spontaneously assembled into small oligomers forming a first β-sheet. Once the β-sheet reached a size of 5-6 monomers, other monomers could interact with the β-sheet surface triggering the formation of a second β-sheet docked by sidechain/sidechain interactions. Once a primary nucleus was formed, we could follow its growth. Each β-sheet provides two ends for elongation, so primary nuclei have four of them. We observed elongation with peptides generally docking from the N-terminus towards the C-terminus as shown in **Figure 6A**. A cross-β protofilament, generally highly twisted, exposes sidechains and termini for secondary nucleation, but we observed that only the sidechains faces could trigger the formation of further β-sheets thus forming a filament (cf. **Figure 6B**). Of note, the addition of each β-sheet decreased the twist. Filaments, made of at least 4 β-sheets, could further growth both through their sidechains faces (cf. **Figure 6B**) as well as through their termini, as exemplified in **Figure 6C**. Growth could occur by N- to N-terminus (head-to-head) as well as N- to C-terminus (head-to-tail) interactions. Importantly, a newly N- to C-β-sheet could grow into a new protofilament, while a newly formed N- to N-β-sheet needed first to shift into an N- to C-one before further growth could occur. A fibril is thus formed when two filaments are linked head-to-tail. Remarkably, the formation of new β-sheets always happened with monomers sliding on the surface before eventually docking. In none of the simulations we observed fragmentation events. Eventually, at high monomer concentration, we also observed interactions between fibrils.

**Figure 6.**
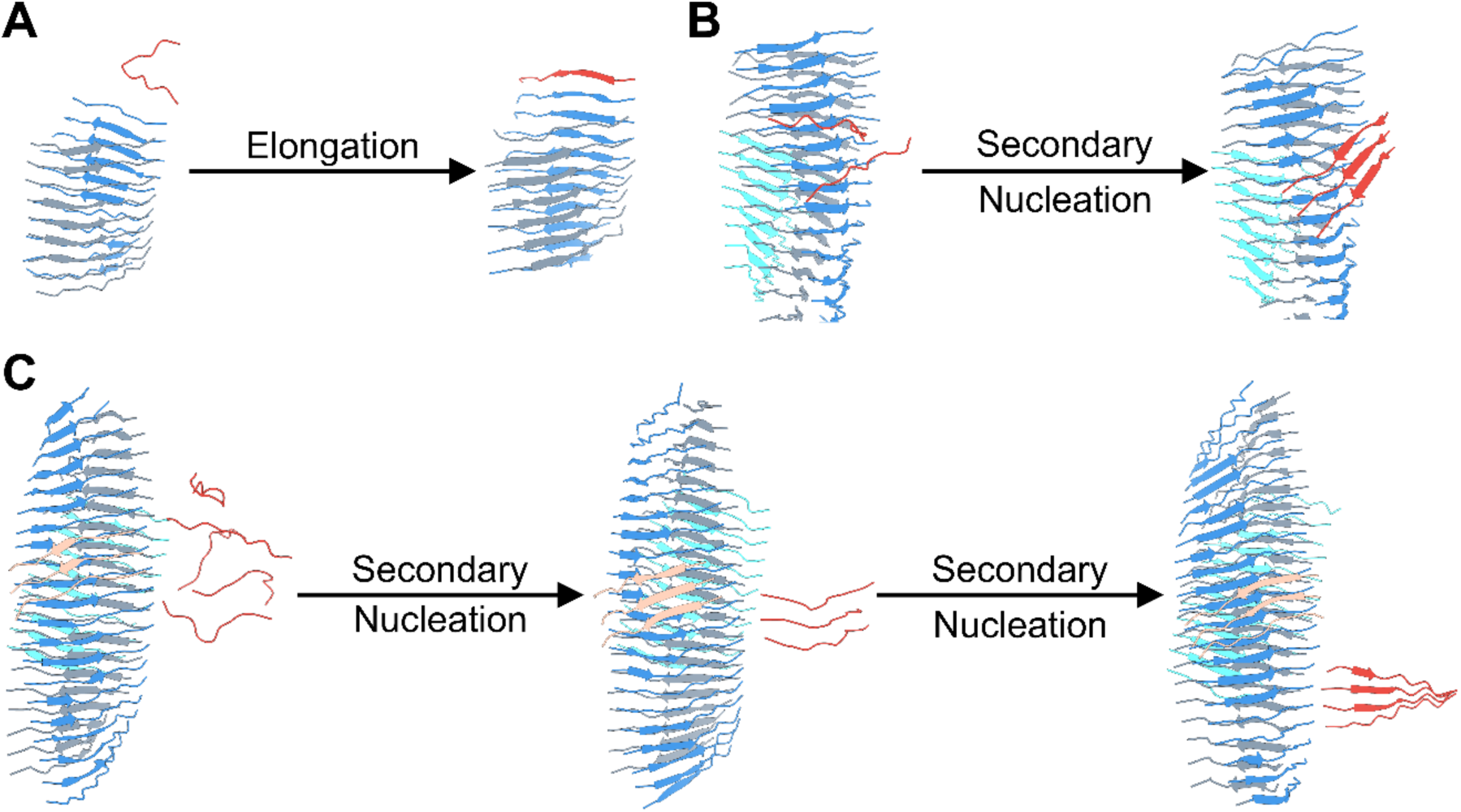
Fibril growth secondary processes. (A) Elongation: the docking of a single monomer in red at one end of a β-sheet in blue. (B) and (C) Surface induced secondary nucleation can happen both on the front surface of a β-sheet (B) as well as on the side surface (C). Peptides can slide on the surface before locking. At least three peptides are required to form a secondary nucleus.

To test if the mechanism described is consistent with the macroscopic kinetics of **Figure 2**, we performed a chemical kinetics analysis of our simulated data using Amylofit (50–52). Simulations could only be fitted globally using a “multi-step secondary nucleation, unseeded model” as shown in **Figure S7**. This is compatible with the positive curvature with an increased slope at lower concentration displayed by *τ*_1/2_ in double log plot (cf. **Figure 2B**) that can be interpreted as the saturation of secondary nucleation (47) at high monomer concentration (e.g., all the catalytic fibril surface is occupied by monomers). Furthermore, our seeded simulations did not show variations in the rate constant *r* (cf. **Figure 2C**), supporting the hypothesis that secondary nucleation is a multi-step process with a first step (monomer attachment on the surface) that is concentration dependent and a second step (monomers rearrangement on the surface) that is concentration independent (47). Amylofit analysis correlates with our observations where the addition of molecules on the cross-β surface implies the exploration of different conformations prior to lock and start forming a new oligomer. Globally, the TTR_105-115_ aggregation process described by our simulations is consistent with the hierarchy proposed by Fitzpatrick et al. (20).

## Discussion

Amyloidogenesis, being the result of an out-of-equilibrium, concentration dependent process, cannot be easily followed by high-resolution structural biology techniques (6). Indeed, while NMR, X-ray crystallography and Cryo-EM have been instrumental to investigate at high resolution both the early steps associated with protein misfolding, as well as the resulting amyloid fibril structures, they have only provided low-resolution information about the transient oligomers populated along the process (53, 54). In this respect, most of the fundamental mechanical understanding of protein aggregation is based on the combination of multiple low-resolution techniques, in particular aggregation kinetics studied by ThT fluorescence, and chemical kinetics analysis (24).

MD simulations could provide the resolution in time and space to observe the emergence of oligomers, nuclei, and fibril from a solution of monomeric proteins (55). This would provide enormous help first to understand the determinants of the different aggregation mechanisms at play, to observe the effect of mutations, and then towards a structure-based design of drugs targeting specific oligomeric species. Unfortunately, computer power is far from being able to enable such simulations using conventional classical mechanics transferable force fields (55). Simulations have thus been employed to study the oligomerization of few peptides (28), the interactions of peptides with preformed fibrils (56), and, in most cases, to understand the determinants of protein aggregation by studying protein monomers only (57–62). These studies, while valuable, fall short in providing indications about oligomers that are by their nature extremely unlikely and short-lived.

Here we set out to develop a simplified force field that could allow studying protein aggregation of thousands of monomers with nowadays state-of-the-art computing resources. Our force field builds on the success of structure based (Gō) models to study protein folding (33, 63). The multi-*e*GO force field introduced here, is 1) at atomic resolution (excluding hydrogens); 2) locally transferable, with bonded and excluded volume interactions derived from a transferable force field or optimized consequently; 3) structure (or ensemble) based using multiple references and symmetrized so to be able to form interactions both intramolecularly as well as intermolecularly. We have shown, in **Figure 1**, that this combination allowed us to describe the conformational ensemble of a disordered peptide in qualitative agreement with more accurate conventional explicit solvent MD simulations. Most importantly multi-*e*GO can describe the aggregation kinetics of thousands of monomers showing concentration dependent features compatible with experiments, **Figure 2-4**. Indeed, the comparison of the scaling exponents *γ* derived from experimental and simulated *τ*_1/2_ data (**Figure 2B** and **Figure 4B**) showed comparable values, confirming the robustness of the model, with a difference observed only for the lowest concentration. TEM morphological analysis of the fibrils highlighted a remarkable degree of polymorphism. Specifically, we classified six main fibril morphologies with highly variable crossover and width (**Figure 4F** and **Table S2**). These data indicate that each fibril is formed by 3 to 8 filaments, in agreement with what observed in *in-silico* fibrils (**Figure 3, Table 1, Table S1** and **S2**). On the contrary all morphologies share a mean width at the crossover of 37.3±4.2 Å, compatible with previous measure, associated with a hydrated cavity that is not observed *in silico* (20). Overall, our model can qualitatively reproduce most of the kinetics and structural features of TTR_105-115_ aggregation, indicating that future improvements should try to better account for solvation effects.

The simulations can then be used to make hypothesis about the oligomeric species populated along the process and provide a structural model for the mechanisms of primary and secondary nucleation. Interestingly, we can show how primary nuclei are in the order of 10 monomers and organized in two opposed β-sheets, **Figure 5D**. These nuclei are populated for less than 0.1% in the lag phase of the kinetics, in comparison to dimers and trimers that are populated around 10 and 1%, respectively (cf. **Figures 5A-C**). This indicates how relevant is to simulate aggregation using large numbers of monomers. Oligomers population drops immediately after the formation of the first nuclei. Following the growth of primary nuclei, our model also shows that elongation happens through the preferential binding of the N-terminus region of the peptide, **Figure 6**. Once a protofilament is formed, secondary nucleation can be observed. Secondary nucleation happens first by the formation of nuclei on the exposed sidechains surface of the filament, then monomers can slide over the surface and eventually dock into it and start the formation of an additional β-sheet layer, **Figure 6**. Once at least four β-sheet layers are formed, we observe additional secondary nucleation events catalyzed by interactions with the N- and C-termini, **Figure 6**. Here, secondary nucleation contributes to the overall growth of fibril while it does not show the formation of independent oligomers that detach to form new protofibrils. We hypothesize that this is a size effect resulting from the relatively small number of monomers that is immediately depleted by the formation of the fibril. Nonetheless the secondary nucleation observed here is still the main effect that contributes to the exponential growth of the fibril mass. Of note, the mechanisms inferred by applying a chemical kinetics analysis on the simulated aggregation kinetics agrees with what was observed in the simulation, **Figure S7**, suggesting that multi-*e*GO could complement and integrate experimental chemical kinetics models to provide a high-resolution time-resolved description of the microscopic processes at play during aggregation (24).

In conclusion, we have presented the first development of multi-*e*GO, a novel structure-based model tailored to study amyloid-type protein aggregation. The model is promising in describing at least qualitatively the spontaneous aggregation of monomers into amyloid fibrils as a function of the initial monomer concentration and thus provide a structural picture of the oligomeric species populated and of the associated aggregation mechanisms. We anticipate that our model can benefit from methodologies that allow integrating simulations with the many complementary experimental techniques deployed to study protein aggregation (64–67). Eventually, the computational efficiency of multi-*e*GO, combined with the availability of the structures of amyloid fibrils of proteins in multiple conditions, will allow to improve our understanding of the mechanisms and the associated oligomeric structures at play in different pathogenic and non-pathogenic self-assembly processes.

## Materials and Methods

MD simulations in this work were performed with GROMACS (68). Models’ parameterization and preparation was developed in python, all scripts, including ad hoc analysis tools, are freely available on GitHub (https://github.com/emalacs/multi-eGO/tree/TTR_paper). Simulations are available on Zenodo (cf. **Dataset S1** and **S2** in the Supporting Information).

### Multi-GO: A multi-reference Gō-like model for protein aggregation

Multi-GO is a multi-reference structure-based force field, at all heavy atom (non-hydrogens) resolution, defined as a combination of terms obtained from two reference structures, namely the protein in its native monomeric state and the amyloid fibril. This was originally developed to study metamorphic proteins (38, 39) using the SMOG software (69). In this model, distances between covalently bonded atoms, as well as angles formed by three subsequent covalently bonded atoms are derived only from the monomeric structure because they describe the local geometry that is generally independent from the specific configuration. Dihedral angles are defined as in SMOG but obtained from both structures and halving the force constant to account for the double counting. Native pairs are obtained from both reference structures following SMOG rules; if two atoms are in contact in both structures, then the distance is defined as the minimum distance. All native pairs are symmetrized so that if atom *i* and atom *j* are in contact in one reference structure, they can interact irrespectively of whether the two atoms belong to the same monomer or to two different monomers; such approach has been successfully employed to describe domain swapped dimers (70, 71), and is needed to make intra- and inter-molecular interactions indistinguishable.

### Multi-eGO: an enhanced Gō-like model for protein aggregation

In contrast to multi-GO, multi-*e*GO force-field is partitioned so that while non-bonded interactions are structure-based, bonded interactions are instead transferable. The multi-*e*GO Hamiltonian given a reference monomer *X*_*m*_ and a reference amyloid structure *X*_*a*_, is defined as:

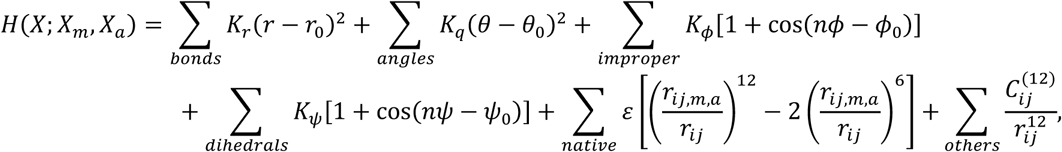

where the parameters for bonds, angles, improper and proper dihedrals are obtained from a transferable force-field, specifically GROMOS54A7 (72) that is a united atom force-field already optimized without non-polar hydrogens. Thus, the local geometry no longer depends on the monomeric structure as in multi-GO. Proper dihedrals terms describing the ϕ and ψ backbone angles were re-optimized as describe in the next section. Interactions between native pairs are defined for couple of atoms farther than one residue (if they belong to the same molecule) and closer than 5.5 Å in either the native monomeric or amyloid structure. In the case of the monomeric state, pairs are obtained from a MD simulation as those forming a contact with a population *P* larger than a threshold *P*_*threshold*_. The interaction length is then defined as the average contact length, while the interaction strength, ε_n_, is heuristically rescaled with respect to a reference ε value as:

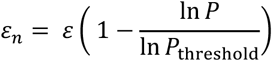

this approach has the merit to give an interaction strength equal to ε if the contact population is 1, like in a static structure, and to smoothly go to 0 when reaching the chosen threshold.

As above all native pairs were symmetrized and if two atoms are in contact in both reference structures, then only the shorter *r*_*ij*_ distance is retained. Care must be taken so that if a native interaction is defined between two atoms belonging to the same or the neighbor residue, this interaction applies only intermolecularly. Finally, excluded volume interactions for all other pairs, that is all pairs of atoms that are not native and that are separated by more than 3 consecutive covalent bonds (*i-i+4*), are defined as in the GROMOS54A7 force field. Among the excluded volume interactions, we include all *i-i+3* interactions involving a backbone nitrogen, in this case the *C*^*(12)*^ Lennard-Jones parameter is scaled down by a factor 0.15. This is needed to effectively account for the missing amide hydrogen, and it is critical to avoid non-physical configurations. Of note, in multi-*e*GO masses are correctly set to include hydrogens; also, since force constants obtained from GROMOS are tuned to work at room temperature, i.e., ∼300 K, *ε*, the reference interaction strength between all native pairs, is a free parameter to be set in a system dependent manner.

### Multi-eGO backbone dihedrals optimization

In the multi-*e*GO Hamiltonian, the intramolecular interactions between atoms belonging to consecutive amino acids are described only by transferable terms. Therefore, a dipeptide simulated with multi-*e*GO should closely mimic the conformational freedom of a dipeptide simulated at room temperature using a transferable force field in explicit solvent. Due to the structure-based non bonded addition, parameters for the proper dihedral angles have been optimized building on the former hypothesis. Alanine, glycine, and proline dipeptides were simulated using CHARMM22* (73) and TIP3P (74) at 300 K for 1 µs each and the resulting Ramachandran distribution was set as our target.

The same dipeptides were simulated using multi-*e*GO, initially setting the force constant *K* of the potential *V*_*D*_ describing proper dihedrals for the backbone ϕ and ψ angles to zero. *V*_*D*_ is defined as *V*_*D*_(*ϑ*) = *K*(1 + cos(*nϑ* − *ϕ*_0_)), with force constant *K*, multiplicity *n*, and phase *ϕ*_*0*._

Target and multi-*e*GO Ramachandran distribution are then compared calculating the following scoring function *S* (i.e., the cross-entropy):

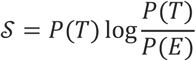

where *P(E)* and *P(T)* are the multi-*e*GO and target Ramachandran distributions. To optimize the parameters *K, n*, and *ϕ*_*0*_ for the proper dihedral angles, we then followed an iterative procedure combining a Monte-Carlo (MC) optimization followed by MD (75, 76). In detail, the effect of a given choice of the parameters is estimated by analytically reweighting the last multi-*e*GO MD simulations as:

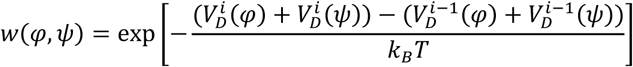

where *V*^*i*^_*D*_ is the potential energy from the *i*th iteration written as the sum of multiple proper dihedral terms; *k*_*B*_ is the Boltzmann constant and *T* the temperature of the MD simulation. Optimal parameters are searched by MC under the constraint that the effective information *N*_*eff*_, calculated over the *N* configurations generated by the last MD, as

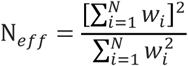

is greater than 0.6. This allows to choose parameters that do not alter dramatically the starting distribution. A new MD simulation is then performed with the chosen parameters and the procedure is repeated till convergence of the scoring function (75, 76).

### Simulations details: TTR_105-115_ monomer in explicit solvent

TTR_105-115_ peptide in solution does not possess a unique well-defined structure being too short, consequently we performed a reference simulation using the AMBER99SBDisp (45) force field. The starting structure was obtained from pdb 4TLT and the simulation was set to resemble the experimental conditions of TTR_105-115_ aggregation, therefore, to be at pH 2.0 we protonated the N- and C-termini. The molecule was solvated with 3408 water molecules in a dodecahedron box initially 0.9 nm larger than the chain in each direction. One chlorine ion was added to neutralize the charges. Short range interactions were cut-off at 1 nm, with long range electrostatic handled using the particle mesh Ewald scheme (77). Lincs constraints were applied only to bonded hydrogens (78).

Following energy minimization, a positional restraint step was performed under NVT conditions at 300K temperature for 500 ps using the velocity-rescale thermostat (79); then the cell-rescale barostat (80) was used to equilibrate the system in the NPT ensemble to the target pressure of 1 atm for 1000 ps. A production MD simulation under NPT ensemble was run for 1.5 µs. This reference simulation was then analyzed to obtain the monomer native pairs and their corresponding interaction strength. These were defined as all the couples of atoms, distant more than one residue, forming a contact with a population *P* larger than a threshold *P*_*threshold*_, 0.09 in this case.

### Simulations details: Multi-eGO TTR_105-115_ fibril

The strength of the native interactions ε is the only parameter to be set empirically based on the knowledge of the system. To find an optimal value for TTR_105-115_, we performed multiple simulations at fixed temperature using a pre-formed fibril. The fibril model was obtained extracting 64 chains from pdb 2M5K (20). Multiple simulations were performed testing a range of ε values between 0.265 and 0.295 kJ/mol at 315 K using a Langevin dynamics with an inverse friction constant of 50 ps and a timestep of 5 fs. The structure was placed in a cubic box 10 nm larger than the fibril in each direction. After the energy minimization and a positional restraint simulation of 1 ns, production simulations were performed for 200 ns. The optimal ε value was chosen to be the highest at which the structure of the fibril was stable with fibril ends monomers showing some flexibility and ability to partially detach from the fibril. An ε value of 0.275 kJ/mol has then been used for all subsequent simulations.

### Simulations details: Multi-eGO TTR_105-115_ aggregation kinetics

To setup the simulations for aggregation kinetics a monomer model was extracted from Y105 to S115 using pdb 4TLT (81) as reference and then, the C-terminal oxygen was added. Four thousand molecules were randomly placed in a cubic box whose side length depends on the desired concentration as *len*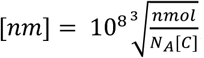, where *nmol* is the number of molecules to add in the system, *N*_*A*_ is the Avogadro Number, and [C] is the concentration. The resulting volumes are in the range of 1 attoliter. The system energy was minimized and then thermalized at a constant temperature of 310 K using positions restraint for 1 ns. Simulations were then performed at 5 different concentration, 13 mM, 11.5 mM, 10 mM, 8.5 mM, and 7 mM in triplicate and evolved long enough to form stable fibrils including most of the monomers. Furthermore, three seeded simulations were performed at 7 mM using a structure of a 10 molecules protofilament obtained at 13 mM as a seed. To analyze the simulations, we used a modified version of the GROMACS *clustsize* tool and homemade python scripts based on MDTraj (82) and MDAnalysis (83, 84). *clustsize* provides a matrix of clusters-sizes as a function of time. The kinetic curves were built by multiplying the number of clusters at every frame by the cluster dimension.

### TTR_105-115_ peptide synthesis

TTR_105-115_ peptide (YTIAALLSPYS) was prepared by microwave assisted Fmoc solid-phase peptide synthesis on Wang resin (0.4 mmol/g loading) using the CEM Liberty Blue synthesizer. The coupling reaction was performed with 5 eq excess of the amino acid (0.2 M in DMF), DIC (0.5 M in DMF) and Oxyma Pure (1M in DMF) as coupling reagents. The MW cycle was as following: 15s at 75°C, 170W followed by 110s at 90°C, 40W. N-Fmoc deprotection was performed using 20% piperidine in DMF with a MW cycle of 15s at 75°C, 155 W, followed by 60 s at 90°C, 50 W. Full-cleavage from the resin was performed by shaking the resin for 3 hours in a mixture of TFA/TIPS/H2O/phenol 90:2.5:2.5:5. After cleavage, the peptide was precipitated and washed using ice-cold diethyl ether. TTR_105-115_ was purified in reverse-phase by using RP-HPLC with a ADAMAS C-18 column from Sepachrom (10 μm, 250 × 21.2 mm, phase A 0.1% TFA in water, phase B: 1% TFA in ACN with a gradient of 20-100% phase B over 40 min at a flow rate of 20 mL/min.).

### TTR_105-115_ aggregation assays

Lyophilized TTR_105-115_ peptide was dissolved at 13 mM in 10% acetonitrile/water solution and pH adjusted to 2.5 with HCl. The solution was sonicated on ice for 15 min and then centrifugated at 4°C for 15 min at 20800 *x g*. The stock solution was eventually diluted to the final concentrations for the experiments (13 mM, 10 mM, and 7 mM) in 10% acetonitrile/water solution. Freshly prepared ThT was added to a final concentration of 20 µM. 50 µL of each condition was then pipetted into black polystyrene 96-well half-area plates with clear bottoms and polyethylene glycol coating (Corning). Each condition was performed in duplicate in each experiment. Plates were sealed to prevent evaporation and incubated at 37 °C under quiescent conditions in a Varioskan Lux plate reader (Thermo Fisher Scientific). Upon excitation at 450 nm, fluorescence at 480 nm was recorded through the bottom of the plate every 120 min. All the experiments were performed in triplicates, except for the 10 mM condition that was done in duplicate. The mean ThT fluorescence values from the independent experiments were normalized and subjected to nonlinear regression analysis, using Boltzmann sigmoidal equation.

### TEM analysis on TTR_105-115_ fibrils

Freshly prepared TTR_105-115_ fibrils were analyzed by TEM. 4-μl droplet of sample was applied onto a 400-mesh copper carbon-coated grids (Agar Scientific) glow discharged for 30 s at 30 mA using a GloQube system (Quorum Technologies). After 1-min incubation, excess of sample was removed and the grid was stained with 2% (wt/v) uranyl acetate solution, blotted dry, and imaged on a Talos L120C transmission electron microscope (Thermo Fisher Scientific) operating at 120 kV. Morphological characterization of the fibrils was performed using the software ImageJ (85).

## Acknowledgments

Funding was provided to C.P. and C.C. by Fondazione Cariplo (CoronAId) and by University of Milano - Linea 1, C.C. is also supported by a grant from Fondazione Telethon (GGP19134). This work was partially supported by Fondazione ARISLA (project TDP-43-STRUCT) and Italian Ministry of Research (PRIN 20207XLJB2) (S.R.). E.S., C.P. and C.C. acknowledge CINECA for an award under the ISCRA initiative, for the availability of high-performance computing resources and support. E.S., C.P. and C.C. acknowledge PRACE for awarding them access to Piz Daint at CSCS, Switzerland. We acknowledge Pietro Sormanni (University of Cambridge), Guido Tiana (University of Milano) and Michele Vendruscolo (University of Cambridge) for their suggestions and insights.

