## Supporting Information for "Multi-*e*GO: an *in-silico* lens to look into protein aggregation kinetics at atomic resolution"

**This PDF file includes:**

Figures S1 to S7
Tables S1 to S2
Legends for Movies S1 to S3
Legends for Datasets S1 to S2

**Other supplementary materials for this manuscript include the following:**

Movies S1 to S3
Datasets S1 to S2

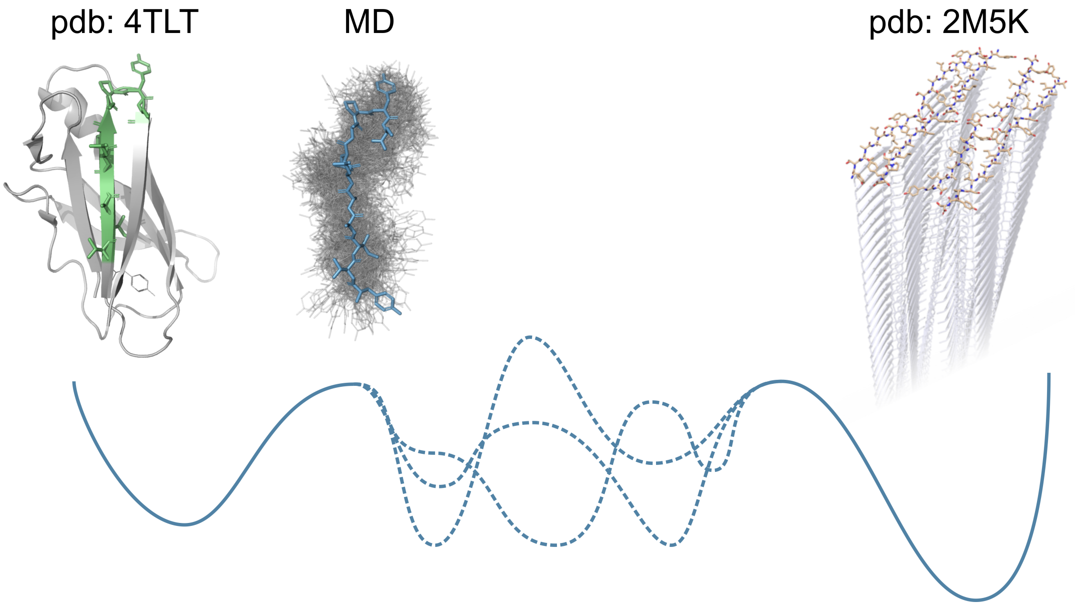


**Figure S1.** Schematic representation of multi-GO and multi-*e*GO models. Multi-GO model is based completely on two structures, one representing the native structure and one the fibril structure. Multi-*e*GO model instead includes information about the conformational dynamics of the native state and the fibril structure, furthermore multi-*e*GO local geometry is described using a transferable potential.

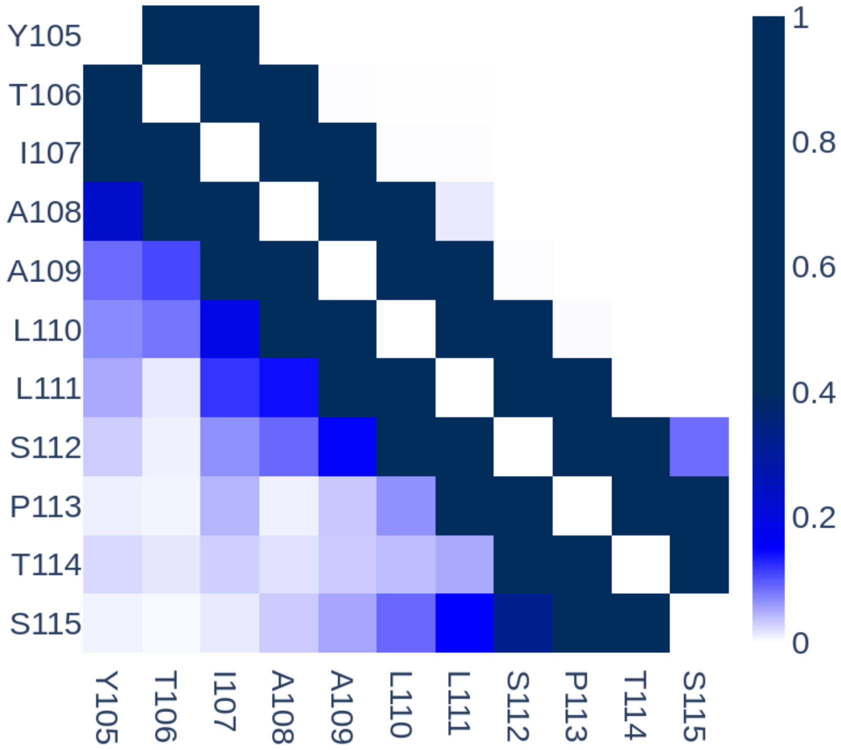


**Figure S2.** Per residue probability contact map for the $TTR_{105\text{-}115}$ peptide conformational ensemble according to a MD simulation with AMBER (lower left) and multi-GO (upper right).

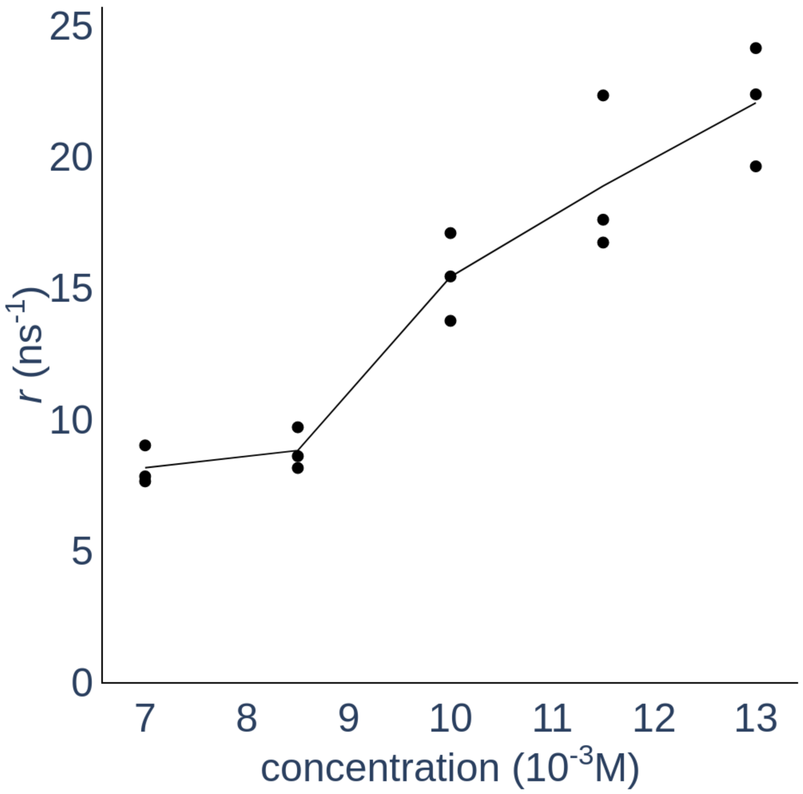


**Figure S3.** Growth rates, *r*, as a function of the initial monomer concentration, for the 15 simulated aggregation kinetics.

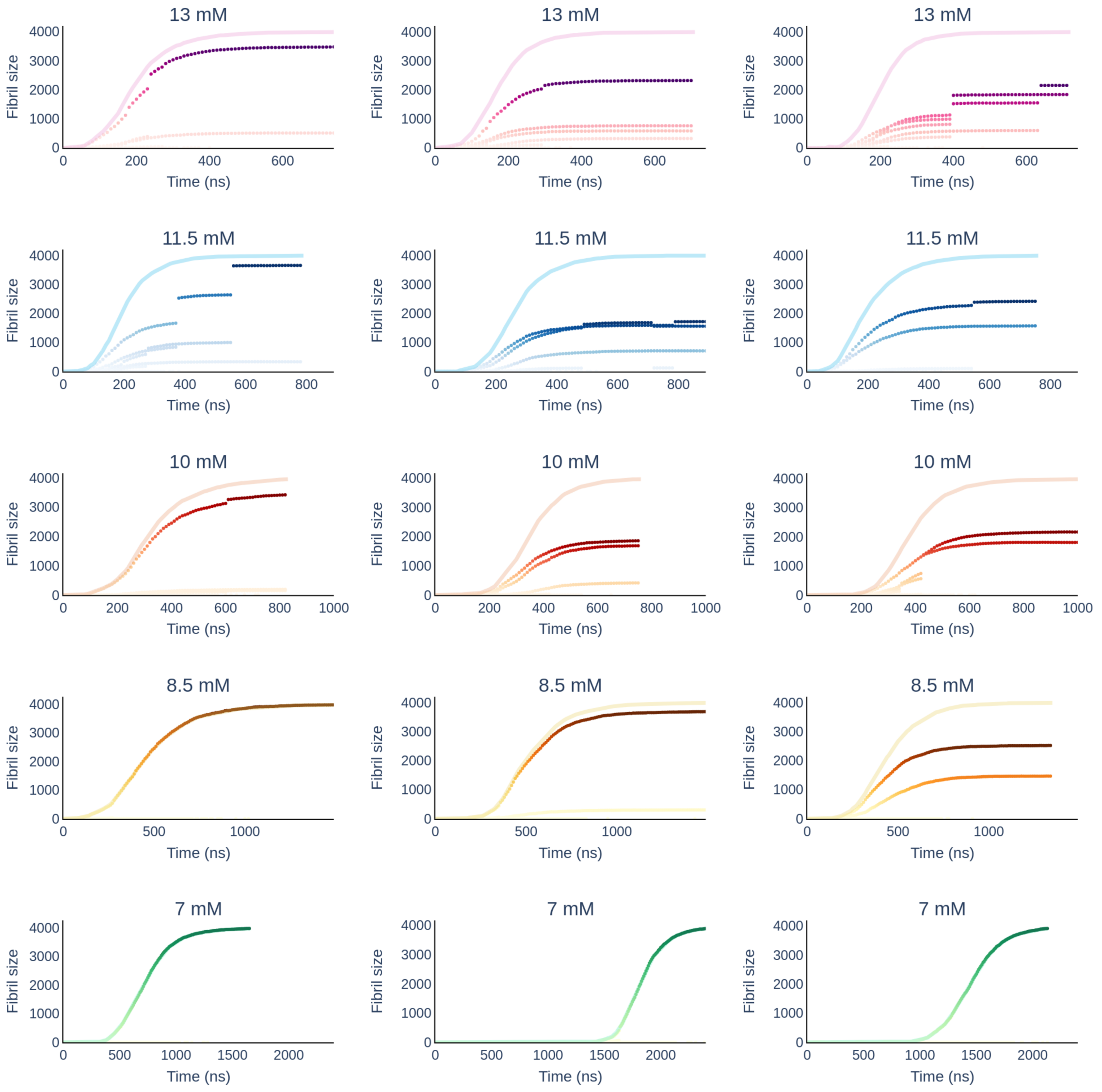


**Figure S4.** Fibrils grow for all simulations. The size of every fibril formed in the simulation is defined by the number of monomers. At high concentration different fibrils are formed and during the simulation they merge into a single fibril as pointed by the gaps.

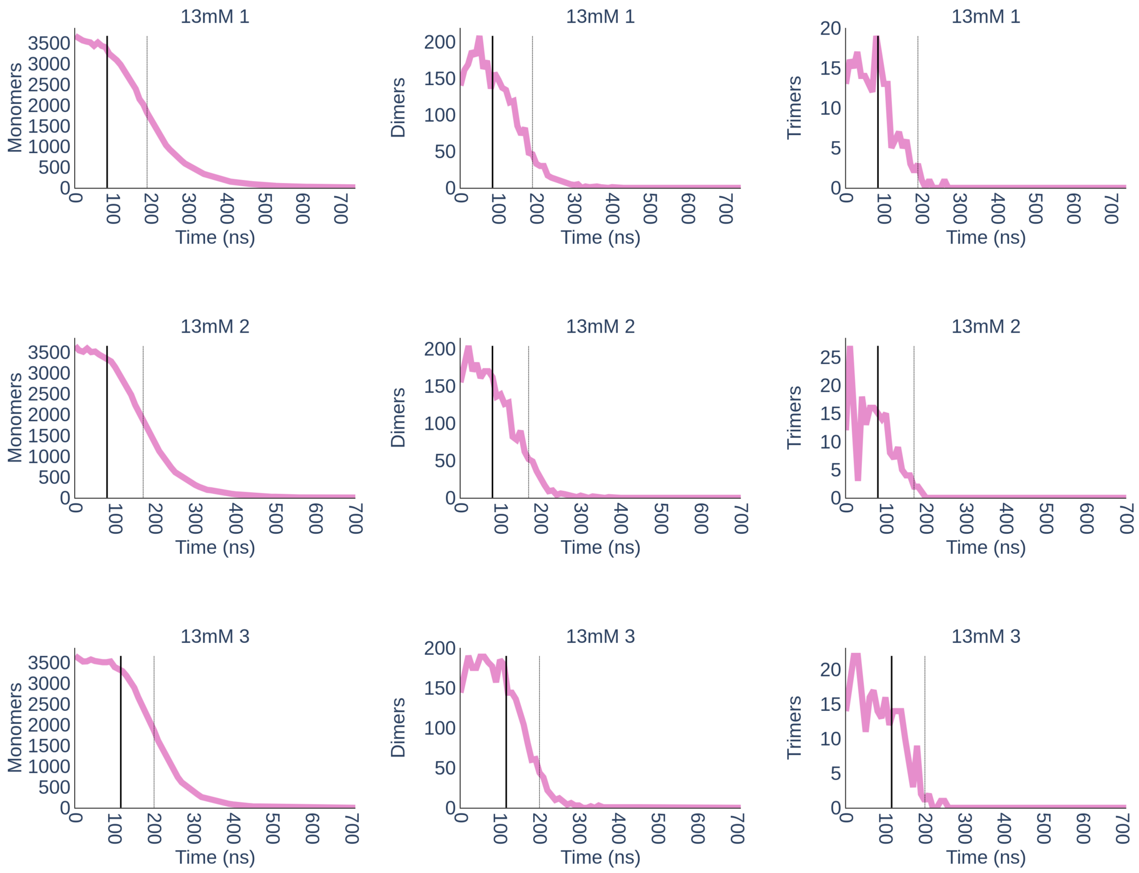
13mM 1
13mM 2
13mM 3
Monomers
Dimers
Trimers
Time (ns)
0
100
200
300
400
500
600
700
500
1000
1500
2000
2500
3000
3500
50
150
5
10
15
20
25

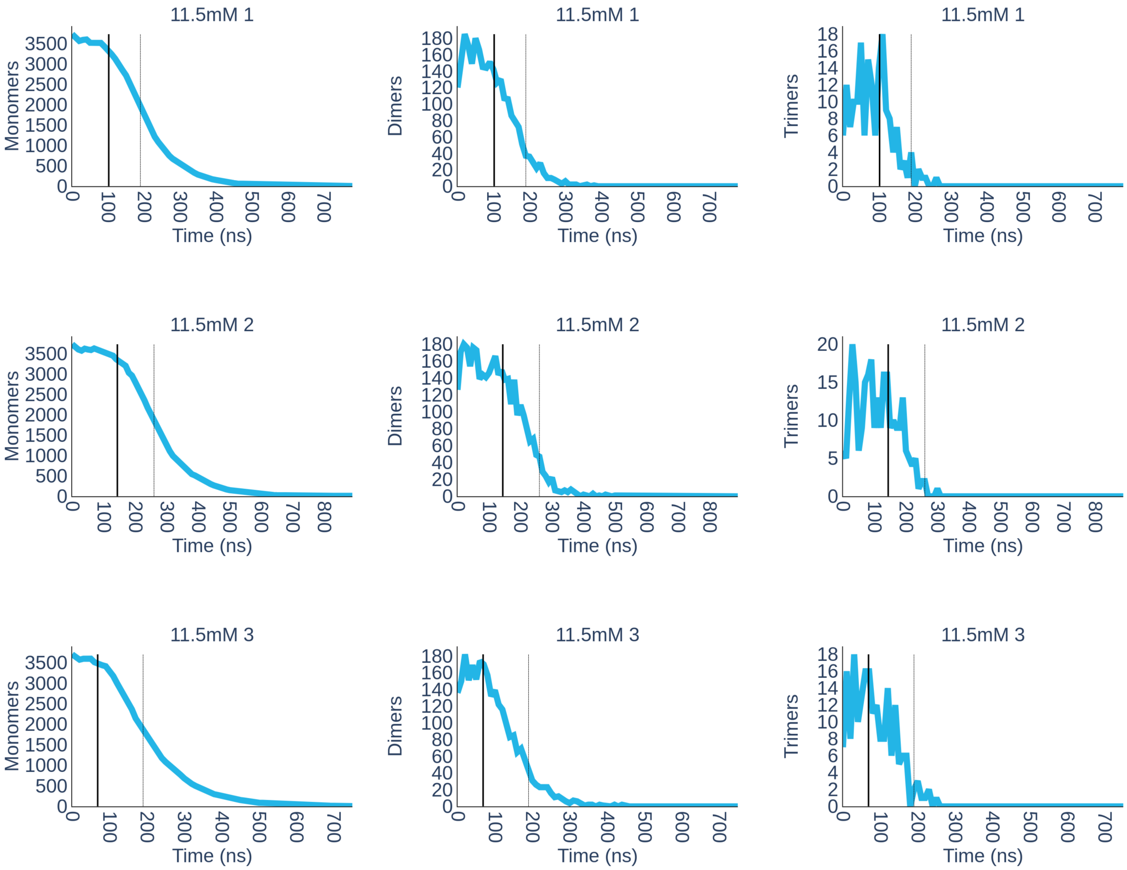

11.5mM 1
Monomers
Time (ns)
11.5mM 1
Dimers
Time (ns)
11.5mM 1
Trimers
Time (ns)
11.5mM 2
Monomers
Time (ns)
11.5mM 2
Dimers
Time (ns)
11.5mM 2
Trimers
Time (ns)
11.5mM 3
Monomers
Time (ns)
11.5mM 3
Dimers
Time (ns)
11.5mM 3
Trimers
Time (ns)

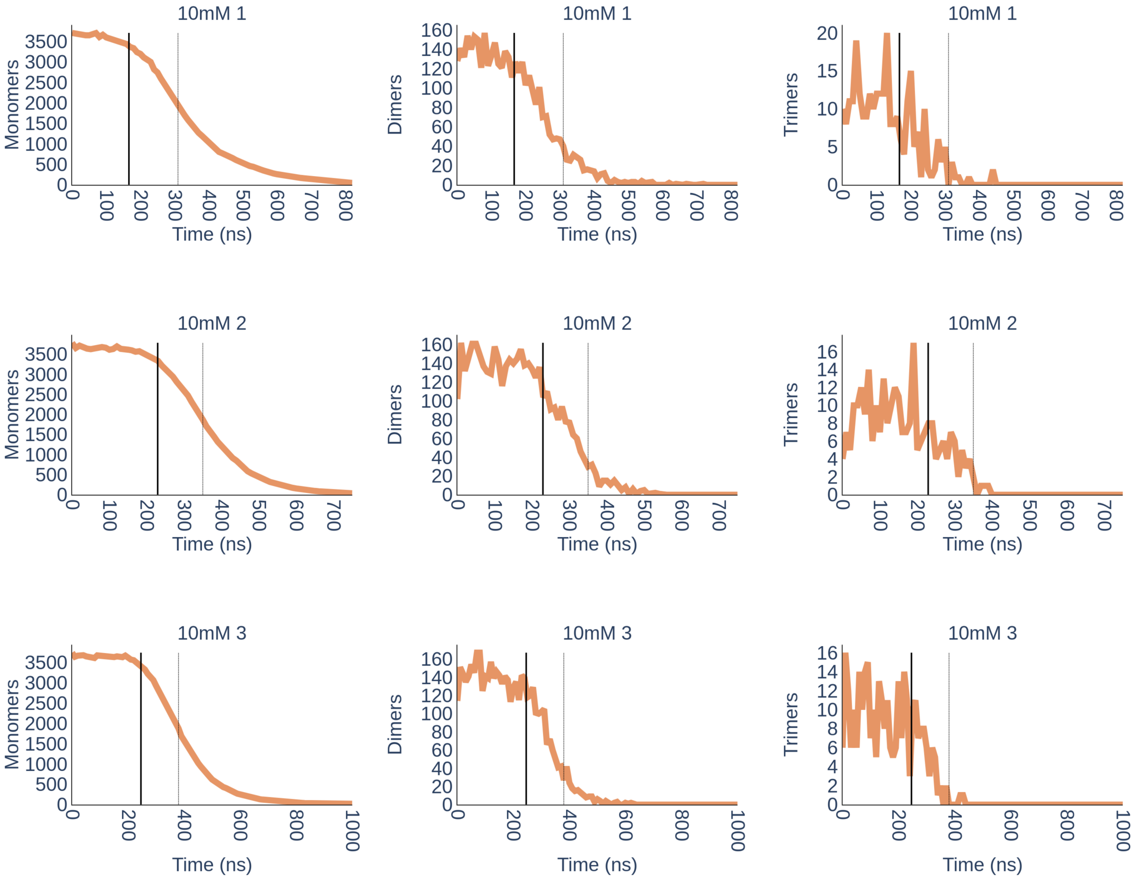
10mM 1
Monomers
Dimers
Trimers
Time (ns)
10mM 2
Monomers
Dimers
Trimers
Time (ns)
10mM 3
Monomers
Dimers
Trimers
Time (ns)

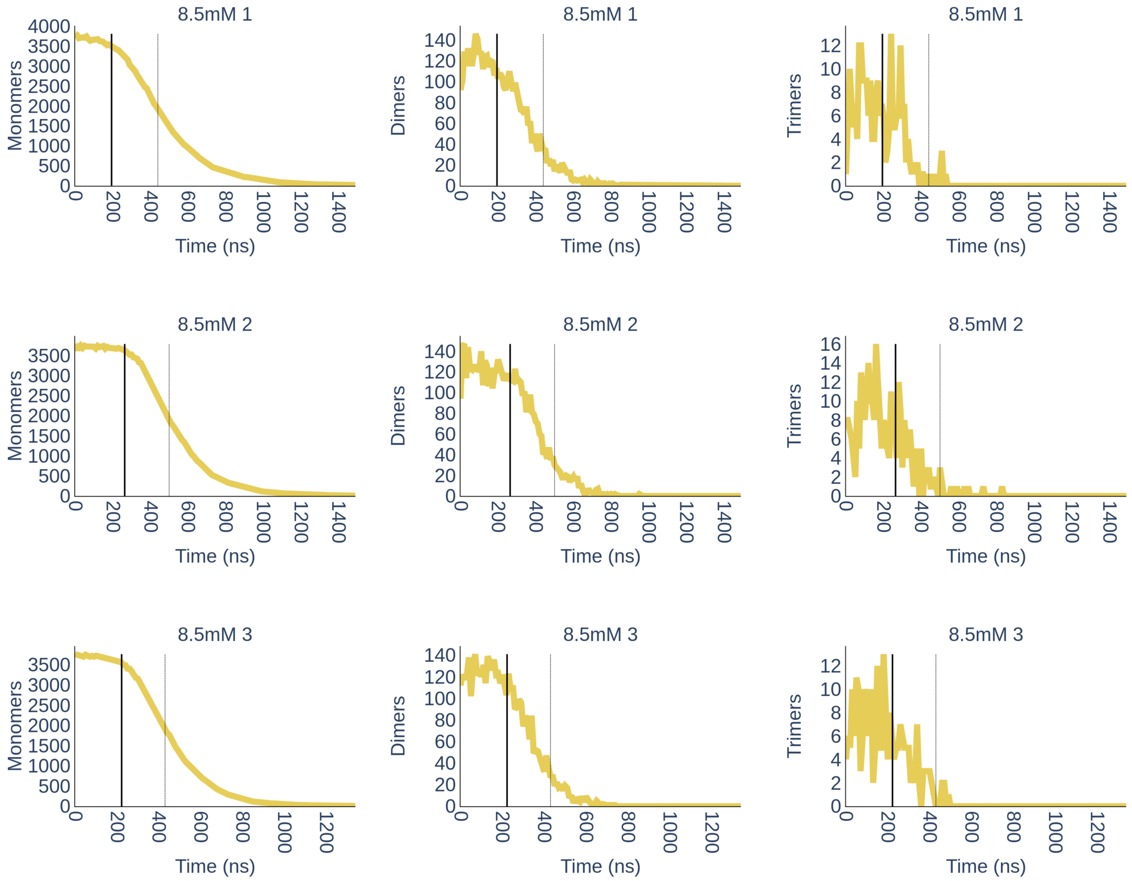
8.5mM 1
Monomers
Dimers
Trimers
Time (ns)
8.5mM 2
8.5mM 3

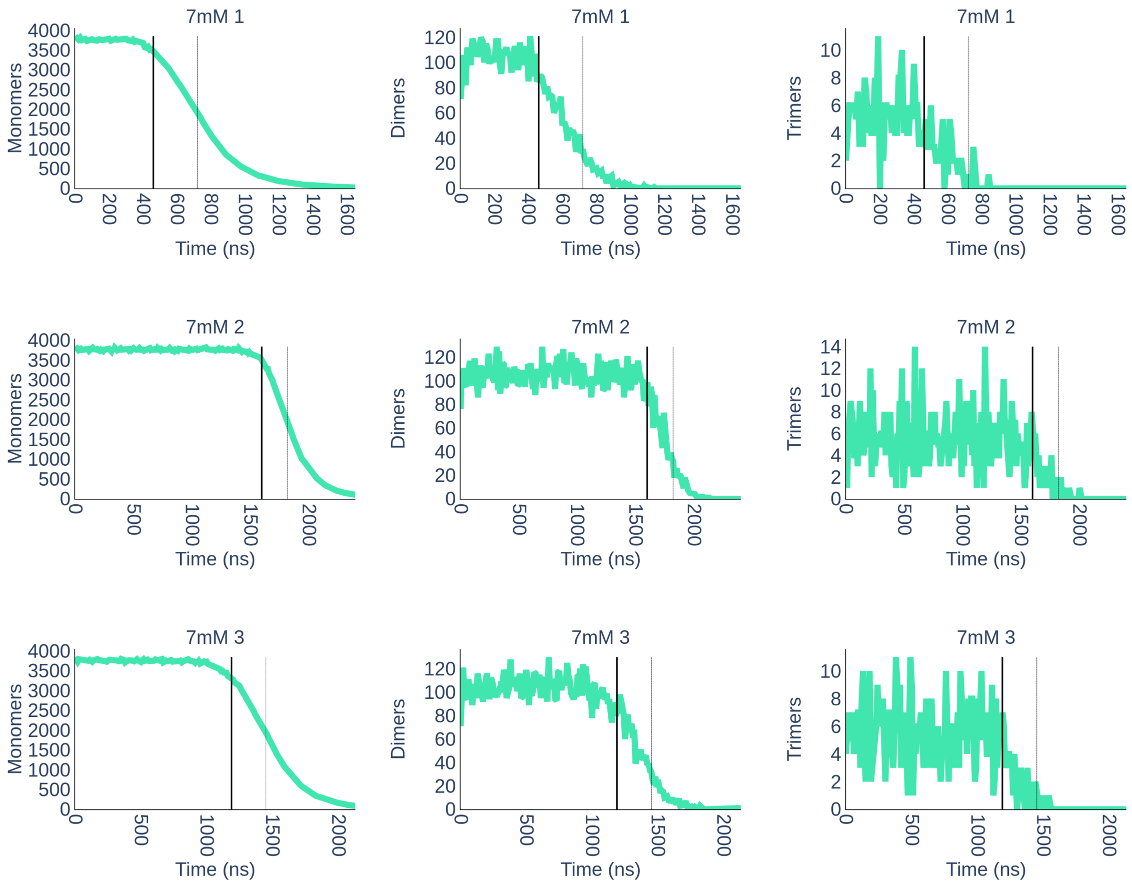


**Figure S5**. Number of monomers, dimers and trimers formed over time for each simulation.

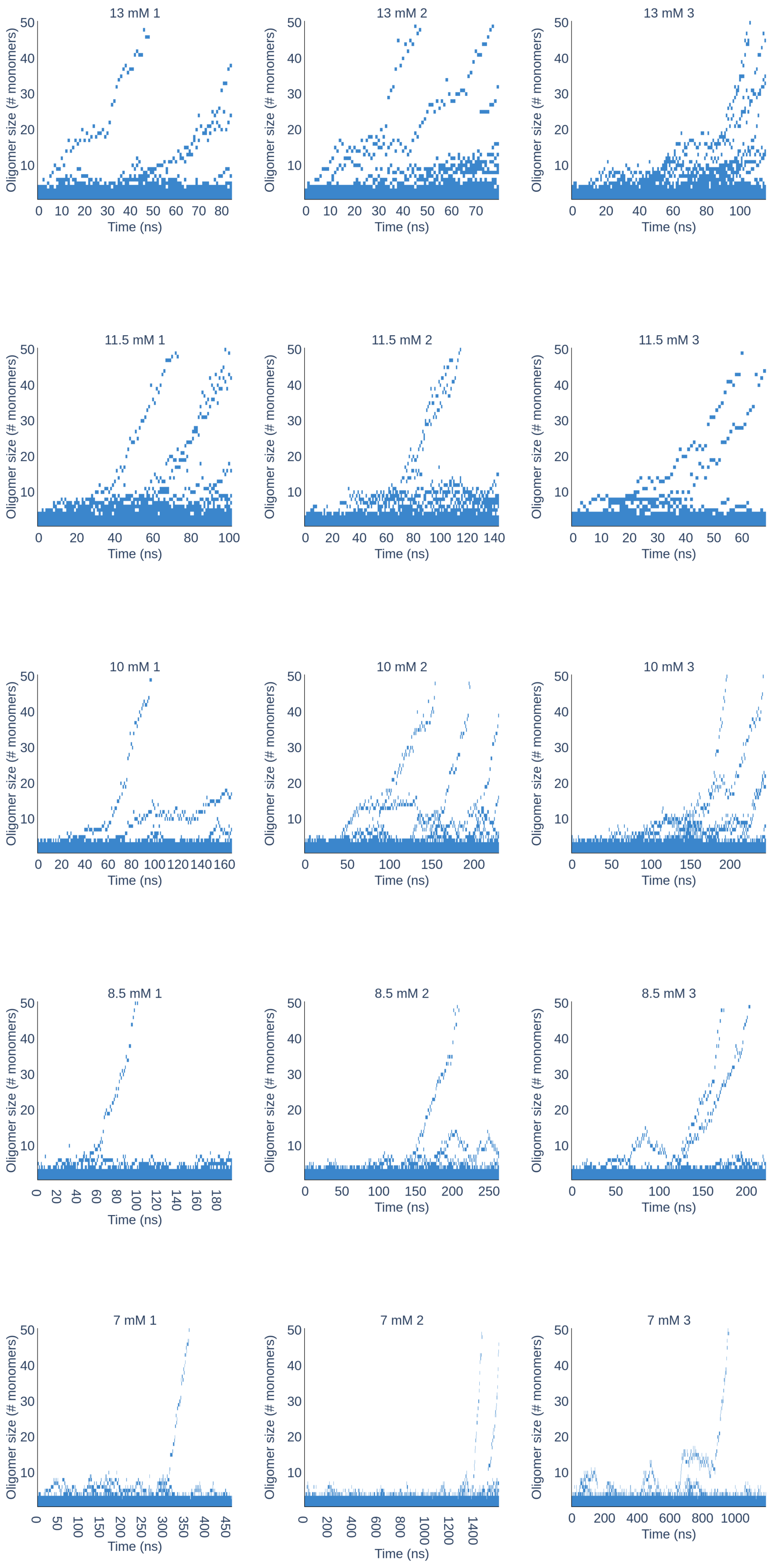


**Figure S6**. Time resolved evolution of oligomers order for all simulations before $t_{lag}$.

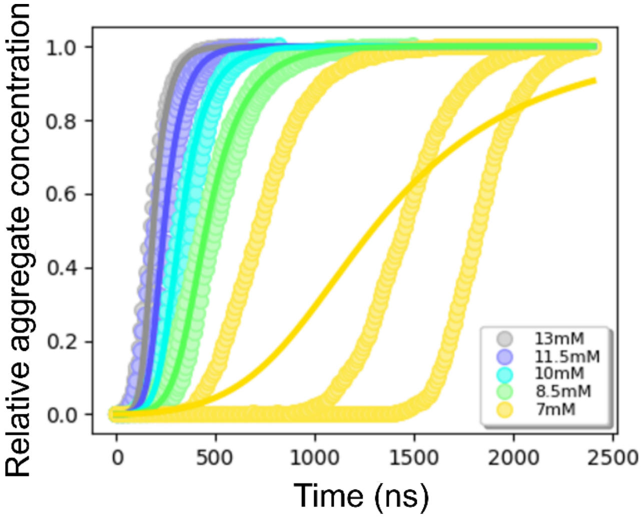


**Figure S7.** Chemical kinetics analysis by Amylofit of the 15 simulated aggregation kinetics traces. The data are fitted using the multi-step secondary nucleation, unseeded model.

**Table S1.** Detailed description of fibrils morphologies obtained *in-silico*.

| | Replica | # Fibrils | Length (Å) | # β-sheets in filaments | # Filaments | Twist (°) | Number of monomers |
|---|---|---|---|---|---|---|---|
| 13 mM | 1 | 4 | 368 | 9 | 5 | -0.22 | 2100 |
| | | | 232 | 9 | 2 | -0.29 | 505 |
| | | | 211 | 5 | 3 | -0.43 | 465 |
| | | | 300 | 7 | 2 | -0.54 | 512 |
| | 2 | 7 | 320 | 10 | 5 | -0.59 | 1513 |
| | | | 198 | 4 | 2 | -0.58 | 206 |
| | | | 175 | 6 | 3 | -0.73 | 336 |
| | | | 100 | 5 | 1 | -4.97 | 86 |
| | | | 246 | 10 | 2 | -0.50 | 324 |
| | | | 221 | 9 | 2 | -0.32 | 584 |
| | | | 163 | 4 | 3 | -0.56 | 324 |
| | 3 | 5 | 380 | 7 | 4 | -0.47 | 1143 |
| | | | 190 | 8 | 2 | -0.38 | 377 |
| | | | 190 | 8 | 3 | -0.18 | 602 |
| | | | 255 | 9 | 4 | -0.33 | 959 |
| | | | 215 | 7 | 4 | -0.5 | 862 |
| 11.5 mM | 1 | 4 | 390 | 6 | 4 | -0.46 | 961 |
| | | | 280 | 12 | 4 | -0.50 | 1717 |
| | | | 220 | 8 | 3 | -0.23 | 909 |
| | | | 210 | 5 | 2 | -0.85 | 336 |
| | 2 | 3 | 330 | 11 | 4 | -0.24 | 1717 |
| | | | 300 | 11 | 3 | -0.24 | 1560 |
| | | | 250 | 5 | 4 | -0.30 | 713 |
| | 3 | 2 | 260 | 13 | 3 | -0.33 | 1574 |
| | | | 310 | 12 | 6 | -0.33 | 2420 |

| | | | | | | | |
|---|---|---|---|---|---|---|---|
| **10 mM** | 1 | 1 | 415 | 9 | 5 | -0.70 | 3425 |
| | 2 | 3 | 366 | 5 | 6 | -0.79 | 1690 |
| | | | 270 | 8 | 4 | -0.23 | 1860 |
| | | | 214 | 5 | 3 | -0.31 | 417 |
| | 3 | 2 | 372 | 9 | 4 | -0.37 | 1811 |
| | | | 315 | 7 | 6 | -0.24 | 2168 |
| **8.5 mM** | 1 | 1 | 515 | 9 | 3 | -0.54 | 3980 |
| | 2 | 1 | 500 | 9 | 4 | -0.15 | 3688 |
| | 3 | 2 | 355 | 14 | 3 | -0.19 | 2524 |
| | | | 370 | 10 | 4 | -0.34 | 1469 |
| **7 mM** | 1 | 1 | 435 | 17 | 5 | -0.11 | 3980 |
| | 2 | 1 | 480 | 7 | 6 | -0.48 | 3359 |
| | 3 | 1 | 490 | 13 | 4 | -0.19 | 3313 |
| **7 mM seeded** | 1 | 1 | 443 | 13 | 5 | -0.35 | 3959 |
| | 2 | 1 | 460 | 14 | 5 | -0.10 | 3907 |
| | 3 | 1 | 485 | 13 | 5 | -0.80 | 3929 |

**Table S2.** Detailed description of *in vitro* fibrils morphologies.

| | Crossover (Å)* | Estimated twist (°) | Maximum width (Å)$^{\Delta}$ | Estimated number of filaments in a fibril• | Width at crossover (Å) |
|---|---|---|---|---|---|
| **I** | 1041 ± 24 | -0.81 | 115 ± 11 | 3 | 39 ± 6 |
| **II** | 1112 ± 57 | -0.76 | 237 ± 28 | 6 | 36 ± 3 |
| **III** | 1121 ± 33 | -0.75 | 171 ± 9 | 5 | 36 ± 2 |
| **IV** | 1185 ± 43 | -0.71 | 326 ± 21 | 8 | 36 ± 4 |
| **V** | 1060 ± 20 | -0.80 | 247 ± 1 | 6 | 35 ± 1 |
| **VI** | 1115 ± 39 | -0.76 | 191 ± 1 | 5 | ND |

* Crossover is defined as the length between two consecutive twists.
˜ Estimated twist is calculated by dividing 180° by the estimated number of molecules in a crossover (calculated considering an average distance between molecules of 4.7 Å).
$^{\Delta}$ Width is defined as the maximum diameter at the largest point within a crossover.
• Calculated considering a peptide length of 38 Å

**Movie S1 (separate file).**
Multi-*e*GO MD trajectory for the aggregation kinetics of the first replica at 13 mM.
Available in Dataset S1.

**Movie S2 (separate file).**
Multi-*e*GO MD trajectory for the aggregation kinetics of the second replica at 7 mM.
Available in Dataset S1.

**Movie S3 (separate file).**
Multi-*e*GO MD trajectory for the aggregation kinetics of the first seeded replica at 7 mM.
Available in Dataset S1.

**Dataset S1 (separate file).**

DOI: 10.5281/zenodo.6125995

Molecular dynamics simulation trajectories of TTR peptide monomers and aggregation kinetics:

- Amber99sb_disp: TTR monomer in explicit solvent using amber99disp force field.
- multi-GO-monomer: TTR monomer simulation using the multi-GO force field
- multi-eGO-monomer: TTR monomer simulation using the multi-eGO ensemble force field.
- multi-eGO-XXmM-Y: aggregation kinetics simulations of TTR using the multi-eGO force field at 13 mM, 11.5 mM and 10 mM concentration; replicate Y
- .ipynb files: analysis scripts employed for the aggregation kinetics simulations.

**Dataset S2 (separate file).**

DOI: 10.5281/zenodo.6125424

Molecular dynamics simulation trajectories of TTR peptide aggregation kinetics:

- multi-eGO-XXmM-Y: aggregation kinetics simulations of TTR using the multi-eGO force field at 8.5 mM, 7 mM and 7 mM seeded; replicate Y.
